# Synapses drive local mitochondrial ATP synthesis to fuel plasticity

**DOI:** 10.1101/2025.04.09.648032

**Authors:** Ilika Ghosh, Ruolin Fan, Monil Shah, Ojasee Bapat, Vidhya Rangaraju

## Abstract

Our brain constantly forms new memories and stabilizes existing memories. To achieve such cognitive flexibility, the brain is wired by plastic synapses that are hotspots of energy consumption. Supplying energy to distant synapses is challenging as they are distributed throughout dendrites and axons, spanning hundreds of microns from their cell body. Synapses, therefore, require an instant and local energy supply provided by mitochondria stabilized near dendritic spines. However, the mechanisms by which synapses communicate their energy demands to locally stable mitochondria to drive local energy production and sustain synaptic plasticity is unknown. Using highly sensitive spine- and mitochondrial ATP reporters and two-photon glutamate uncaging to stimulate individual spines, we find that synaptic plasticity input drives instant and sustained increase in spine ATP levels, provided by local ATP synthesis in ∼10-20 μm spatially confined compartments within mitochondria. This spatially localized mitochondrial ATP generation is driven by a spatially localized mitochondrial calcium influx independent of the endoplasmic reticulum. Notably, the initial spine ATP increase, supported by local mitochondrial ATP synthesis, is independent of CaMKII and the energy demands of spine structural plasticity. Without local calcium signaling and mitochondrial stabilization, synapses do not meet their instant and sustained energy needs, resulting in synaptic plasticity defects, as observed in neurological disorders.

## Introduction

The human brain is unrivaled in its energy production efficiency, precisely adjusting its energy supply in space and time where and when the needs arise. Yet, this evolutionary triumph brings vulnerability; even brief disruptions in brain energy supply lead to severe cognitive deficits^1^. Synapses, the brain’s functional units, are hotspots of energy consumption during activity, given their changing energy demands in maintaining ionic balance, synthesizing proteins, trafficking molecules, and cellular signaling. Supplying the energy currency ATP to synapses distributed throughout the expansive networks of dendrites and axons, spanning distances as long as meters from their cell body, is a monumental task requiring precise delivery tailored to each synapse’s unique activity history.

Consistent with this notion, we recently showed that mitochondria are locally stabilized within dendrites by a vesicle-associated membrane protein-associate protein (VAP) to support synaptic plasticity and the associated synaptic protein synthesis^2,3^. While mitochondrial dysfunction is associated with various neurological disorders such as Alzheimer’s, Parkinson’s, and Amyotrophic Lateral Sclerosis^4,5^, several fundamental mechanisms of dendritic spine energetics and mitochondrial energy supply are unknown, including: (1) what are the absolute steady-state ATP levels of dendritic spines and how are they maintained; (2) what is the impact of a synaptic plasticity input on the ATP levels of a dendritic spine; (3) do mitochondria generate ATP to meet synaptic plasticity’s energy demands, and is it local; (4) what is the molecular signal that drives local mitochondrial ATP generation to support synaptic plasticity; (5) do changes in this local molecular signaling pathway impact synaptic plasticity and disease.

The challenge in understanding spine energetics and local mitochondrial ATP provision primarily arises from the lack of sensitive ATP reporters to measure ATP within tiny compartments of spines and mitochondria. There is also a need to combine spine and mitochondrial ATP measurements with single spine stimulation and plasticity induction. We have developed an imaging approach that uses newly engineered spine- and mitochondria-targeted ATP reporters to combine ATP measurements at single-spine and -mitochondrial resolution upon spine plasticity induction using two-photon glutamate uncaging. We find an instant and sustained increase in spine ATP levels provided by local mitochondrial ATP production, enabled by local mitochondrial stabilization and calcium signaling. Surprisingly, the initial spine ATP burst contributed by mitochondrial ATP synthesis is not driven by energy demand. Without local mitochondrial ATP production, synapses lose their ability to ramp up and sustain ATP levels when needed, affecting synaptic plasticity formation and sustenance, as observed in neurological disorders such as Amyotrophic Lateral Sclerosis (ALS)^2^.

## Results

### A quantitative reporter of Spine ATP

To measure spine ATP levels, we modified our previously developed presynaptic ATP reporter based on an engineered firefly luciferase^6^. Luciferase catalyzes the oxidation of luciferin, a cell-permeant substrate, using ATP and Mg^2+^ to produce light by bioluminescence. To measure ATP within dendritic spines, we fused luciferase (luc) to Homer2, a postsynaptic density scaffolding protein. To calibrate for reporter expression level, we fused luc with a pH-sensitive but thermo- and photostable fluorescent protein mOrange2 (mOrg)^7^, allowing for pH-corrected, ratiometric luminescence to fluorescence (L/F) readouts (see Methods). We refer to this ratiometric spine ATP reporter as Spn-ATP and the spine ATP measurements as ATP_spine_ (Fig. 1A).

**Figure 1.**
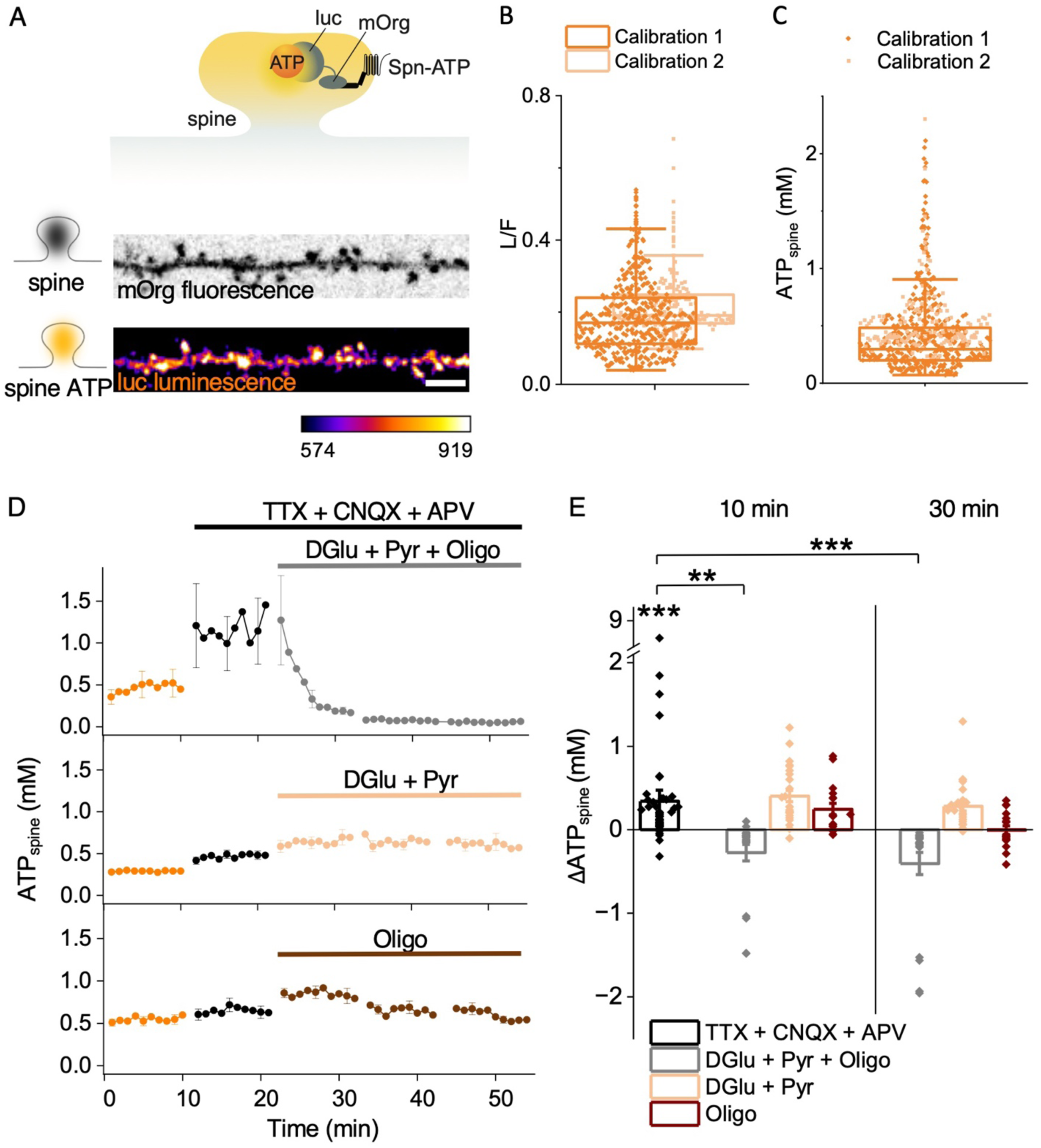
Steady-state ATP_spine_ is heterogenous across spines and is supported by glycolysis and mitochondrial ATP synthesis A (top) Illustration showing the design of the spine ATP sensor, Spn-ATP, for spine ATP measurements. Luciferase (luc) is conjugated to the fluorescent and pH-sensitive protein, mOrange (mOrg), and targeted to the spine via Homer2 (black rectangle). **(bottom)** Representative images (of **B**, **C**) showing mOrg fluorescence (black) and luc luminescence (orange) from Spn-ATP. The intensity scale is in arbitrary units. **B** The average L/F measurements obtained using Calibration 1 (orange) and 2 (light orange) (see Methods) from individual spines show heterogeneity. n in spines, animals: 543, 41. **C** The average spine ATP measurements obtained using Calibration 1 (orange) and 2 (light orange) show a mean of 0.41 ± 0.01 mM with a CV of 82%. n in spines, animals: 543, 41. **D.** The average time course of ATP_spine_ (orange) upon global neuronal activity silencing (black, TTX+CNQX+APV) shows an increase. Upon blocking glycolysis and mitochondrial oxidative phosphorylation (gray, DGlu+Pyr+Oligo), ATP_spine_ plummeted with an exponential time constant of (τ) of 3.0 ± 0.1 min, while negligible change was observed when either glycolysis (light orange, DGlu+Pyr) or mitochondrial oxidative phosphorylation (brown, Oligo) was blocked. n in spines, animals: 23, 2 (DGlu+Pyr+Oligo), 25, 2 (DGlu+Pyr), 17, 2 (Oligo). **E.** The average change in ATP_spine_ (ΔATP_spine_) shows a significant increase upon global neuronal activity silencing (black, TTX+CNQX+APV) compared to baseline ATP_spine_, and a substantial decrease upon glycolytic and mitochondrial ATP synthesis block 10- and 30-min post inhibition (gray, DGlu+Pyr+Oligo) compared to the global neuronal activity silencing baseline (black, TTX+CNQX+APV). n in spines, animals: 23, 2 (DGlu+Pyr+Oligo), 25, 2 (DGlu+Pyr), 17, 2 (Oligo). One-way ANOVA, Tukey test, p-values: 0.0003 (Baseline vs. TTX+CNQX+APV), 0.00713 (10 min post inhibition: TTX+CNQX+APV vs. DGlu+Pyr+Oligo), 0.99977 (10 min post inhibition: TTX+CNQX+APV vs. DGlu+Pyr), 0.99888 (10 min post inhibition: TTX+CNQX+APV vs. Oligo), 0.0004 (30 min post inhibition: TTX+CNQX+APV vs. DGlu+Pyr+Oligo), 0.99979 (30 min post inhibition: TTX+CNQX+APV vs. DGlu+Pyr), 0.54323 (30 min post inhibition: TTX+CNQX+APV vs. Oligo). See also Figure S1.

As Spn-ATP is luminescence-based, it emits significantly fewer photons than fluorescence-based reporters. Therefore, a conventional fluorescence microscope is insufficient to visualize the luminescence signal. Furthermore, to mimic a presynaptic stimulus by local glutamate uncaging to stimulate individual spines and induce synaptic plasticity, the microscope needs an integrated fluorescence imaging system and a two-photon photomanipulation system. Therefore, we custom-built a microscope with two cameras – a highly sensitive camera (EMCCD) to measure luminescence signal and a second camera for confocal fluorescence imaging and two-photon spine stimulation and plasticity induction (see Methods). Expression of Spn-ATP in hippocampal neurons resulted in the efficient delivery of the reporter within dendritic spines, visualized by mOrg fluorescence and luc luminescence (Fig. 1A).

We calibrated the L/F measurements from Spn-ATP to steady-state ATP_spine_ concentrations by obtaining an ATP titration curve using a previously established method^6^ (Fig 1B, C, S1A-C, see Methods). The calibration method was validated by parallel measurements in the soma of the same neurons, confirming that the ATP_soma_ concentrations obtained by this calibration method are comparable to previous observations (1-3 mM)^8–11^ (Fig. S1D). We find that at steady-state, ATP_spine_ measured at a 50 – 100 μm distance from the soma is 0.41 ± 0.01 mM (mean±SEM), averaged over 543 spines from 109 neurons, CV = 82% across spines, 67% across neurons (Fig. 1C, S1C), corresponding to ∼10^5^ molecules in a typical dendritic spine (see Methods). Although Spn-ATP itself consumes ATP, this burden is insignificant as the total flux of photons corresponds to only 5168 ATPs/min/spine, 0.1% of the reported ATP production rate in neurons 6X10^6^ ATPs/min/spine^12^ (see Methods). Interestingly, steady-state spine ATP decreases along the dendritic length with increasing distance from the soma (Fig. S1E), indicating the need for local energy supplies to meet the instant and sustained energy demands of distant spines. No correlation was observed between baseline spine size and ATP_spine_ (Fig. S1F).

### Resting ATP_spine_ is tightly regulated by neuronal and spine activity

One of the fundamental questions about dendritic spine energetics is whether the activity state of the neuron influences resting spine ATP levels. To address this question, we measured resting ATP_spine_ upon silencing global neuronal activity by adding the Na^+^ channel blocker, TTX, to block action potentials and the AMPAR and NMDAR blockers, CNQX and APV, to eliminate any postsynaptic input from spontaneous presynaptic vesicle fusion and neurotransmitter release. Upon acute global neuronal silencing for 10 minutes, ATP_spine_ significantly increased compared to baseline (Fig. 1D, E), suggesting that resting ATP_spine_ is tightly regulated by neuronal and spine activity. To examine the sources supplying energy to resting ATP_spine_, we treated the acute globally silenced neuron with inhibitors that block either or both of the two main ATP synthesis pathways: (1) Glycolysis was inhibited using the competitive inhibitor of glucose-6-phosphate, 2-deoxyglucose, in the presence of the mitochondrial substrate pyruvate to keep mitochondrial oxidative phosphorylation unaffected (DGlu+Pyr)^3^; (2) Mitochondrial oxidative phosphorylation was inhibited using the F1F0-ATP synthase inhibitor, oligomycin (Oligo)^6^. Upon acute treatment of the silenced neurons with both inhibitors for 10 minutes, ATP_spine_ plummeted with an exponential time constant (τ) of 3.0 ± 0.1 min that did not recover for at least 30 minutes post-inhibition, suggesting that even in the absence of action potential and postsynaptic input from spontaneous vesicle fusion, the housekeeping cellular processes contribute to significant energy demand on resting ATP_spine_, fueled by glycolytic and mitochondrial ATP synthesis (Fig. 1D, E). When only glycolysis or oxidative phosphorylation was inhibited, ATP_spine_ remained unchanged, suggesting that at resting state, in the absence of one ATP_spine_ source, the other can compensate (Fig. 1D, E).

### Synaptic plasticity induction increases ATP_spine_

Repetitive activation of a postsynaptic spine causes the potentiation of its synaptic strength, called synaptic plasticity^13^. Upon synaptic plasticity induction, many downstream processes are triggered within the spine that are energy-consuming, such as maintenance of ionic gradients, kinase-dependent protein signaling, actin polymerization-mediated spine size increase, protein synthesis, active transport and vesicular trafficking of key molecular players to the spine^13^. We wanted to determine the impact of synaptic plasticity on spine ATP levels. Using previously established methods, we induced synaptic plasticity by stimulating individual spines with two-photon glutamate uncaging at 0.5 Hz, 60 s^2,3,14^ (Fig. 2A-D). Synaptic plasticity induction was confirmed by the size increase of the plasticity-induced spine, otherwise called spine structural plasticity, using previously established methods^2,3,14^ (Fig. 2E). We expected two scenarios upon synaptic plasticity induction, similar to observations in presynaptic terminals^6,15,16^: (i) spine ATP levels will remain unchanged due to a tight link between energy demand and energy synthesis or (ii) spine ATP levels will decrease in response to the significant energy demands of synaptic plasticity formation and maintenance.

**Figure 2.**
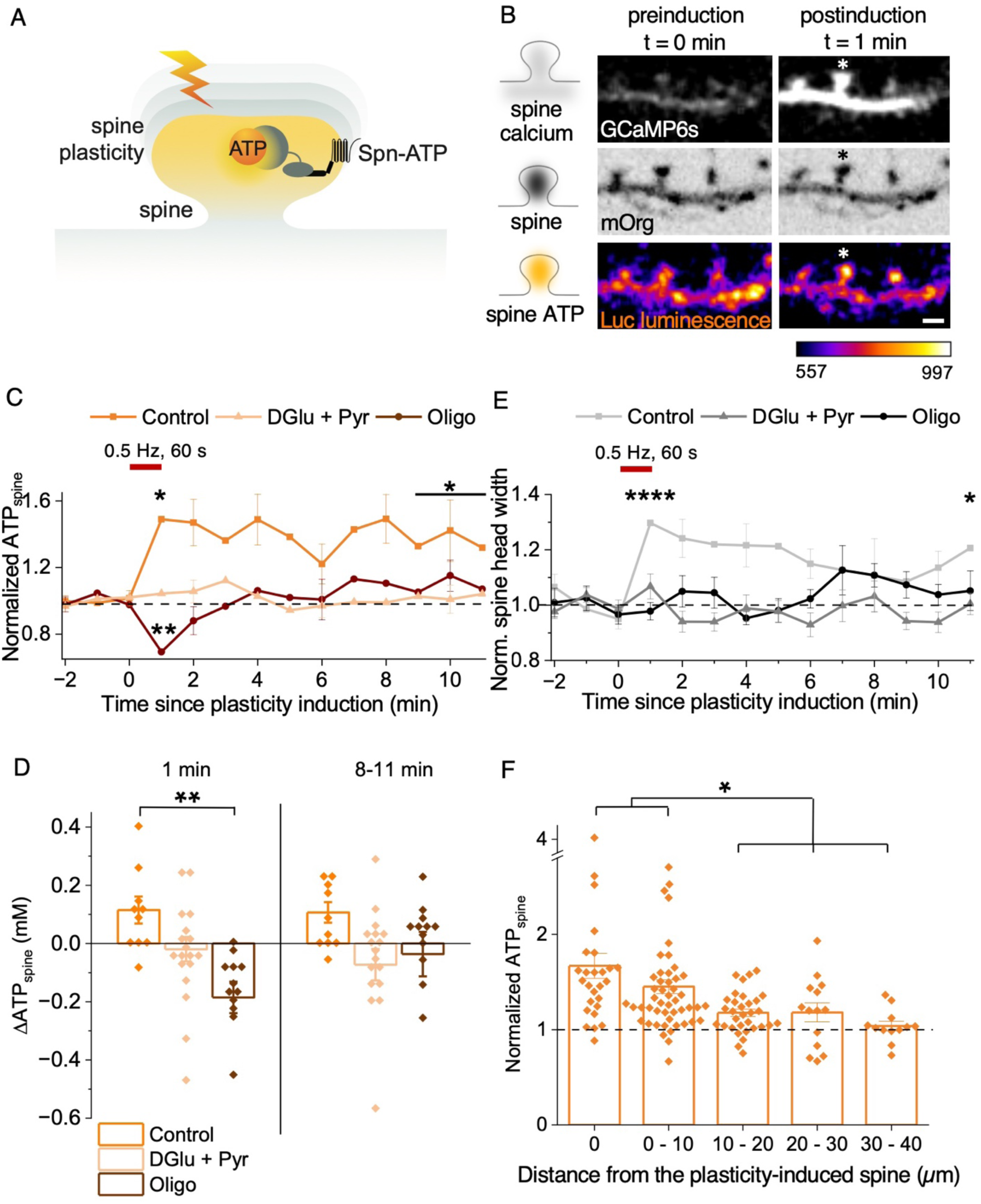
Synaptic plasticity induction increases ATP_spine_ **A** Illustration showing Spn-ATP for ATP_spine_ measurements following synaptic plasticity induction (lightning bolt). **B** Representative images (of **C, D, E**) showing spine calcium (white, GCaMP6s) and Spn-ATP luminescence (orange, luc) increase upon synaptic plasticity induction (white and black asterisk) without a change in spn-ATP expression (black, mOrg). Note that only 1 out of 10 spines show an apparent increase in raw luminescence signal upon synaptic plasticity induction, while in the rest of the spines, the instant increase in spine ATP is only evident after pH correction and conversion of the L/F ratio to ATP_spine_ values using the ATP titration curve (see S2A, B). The intensity scale is in arbitrary units. **C** The average time course of ATP_spine_ upon synaptic plasticity induction (red bar, 0.5 Hz, 60 s) shows an instant increase sustained for up to 11 min post-plasticity induction in Control (orange). The instant and sustained increase is lost upon mitochondrial ATP synthesis inhibition (brown, Oligo), while there was negligible change upon glycolytic inhibition (light orange, DGlu+Pyr). n in spines, animals: 10, 3 (Control), 18, 3 (Dglu+Pyr), 13, 3 (Oligo). Paired sample t test, p-values: 0.03516 (Control: Baseline vs. 1 min post-plasticity induction), 0.01461 (Control: Baseline vs. average of 9-11 min post-plasticity induction), 0.6407 (DGlu+Pyr: Baseline vs. 1 min post-plasticity induction), 0.00509 (Oligo: Baseline vs. 1 min post-plasticity induction). **D** The average change in ATP_spine_ (ΔATP_spine_) shows a significant decrease upon mitochondrial ATP synthesis block (brown, Oligo) 1 min post-plasticity induction compared to the Control (orange) but not upon glycolytic inhibition (light orange, DGlu+Pyr). There was no significant difference 8-11 min post-plasticity induction. n in spines, animals: 10, 3 (Control), 18, 3 (Dglu+Pyr), 13, 3 (Oligo). One-way ANOVA, Tukey test, p-values: 0.54182 (1 min post-plasticity induction: Control vs. DGlu+Pyr), 0.00885 (1 min post-plasticity induction: Control vs. Oligo), 0.22527 (8-11 min post-plasticity induction: Control vs. Dglu+Pyr), 0.54653 (8-11 min post-plasticity induction: Control vs. Oligo). **E** The average time course shows a sustained rise in spine-head width at 1- and 11-min post-plasticity induction (red bar, 0.5 Hz, 60 s). n in spines, animals: 6, 2 (Control), 12, 3 (DGlu+Pyr), 8, 2 (Oligo). Paired sample t test, p-values: <0.0001 (Control: Baseline vs. 1 min post-plasticity induction), 0.04345 (Control: Baseline vs. 11 min post-plasticity induction). One-way ANOVA, Tukey test, p-values: 0.17453 (1 min post-plasticity induction: Control vs. Dglu+Pyr), <0.0001 (1 min post-plasticity induction: Control vs. Oligo), 0.32204 (11 min post-plasticity induction: Control vs. Dglu+Pyr), 0.03531 (11 min post-plasticity induction: Control vs. Oligo). **F** The normalized ATP_spine_ is significantly increased upon synaptic plasticity induction within 0-10 μm, but not within 10-40 μm, from the plasticity-induced spine. n in spines, animals: 27, 4 (0 μm), 50, 4 (0-10 μm), 32, 4 (10-20 μm), 14, 3 (20-30 μm), 12, 3 (30-40 μm). One-way ANOVA, Tukey test, p-values: 0.29114 (0 vs 0-10 μm), 0.000886 (0 vs. 10-20 μm), 0.01596 (0 vs. 20-30 μm), 0.0015 (0 vs. 30-40 μm). See also Figure S2.

Contrary to both expectations, ATP_spine_ increased 1.5-fold instantly 1 min following plasticity induction and remained sustained for at least 11 minutes post-plasticity induction (Fig. 2B-D, S2A, B, Supp. Video 1, see Methods). This increase was not due to the movement or trafficking of Spn-ATP (Fig. S2C, see Methods). We also did not see a strong correlation between baseline spine size and ATP_spine_ increase, baseline ATP_spine_ and ATP_spine_ increase, or spine size increase and ATP_spine_ increase upon synaptic plasticity induction (Fig. S1F). Mitochondria drive the instant and sustained increase 1 min and 11 min post-plasticity induction as inhibition of mitochondrial oxidative phosphorylation (using Oligo) resulted in a significant decrease in ATP_spine,_ which, while recovered to baseline ATP levels, did not recover to the 1.5-fold sustained increase seen in the Control neurons (Fig. 2C, D). Interestingly, glycolytic inhibition (using Dglu+Pyr) also affected the instant and sustained rise in ATP_spine_, compared to the Control, but did not show a significant decrease in ATP_spine_ as seen with inhibition of mitochondrial oxidative phosphorylation (Fig. 2C, D). Spine structural plasticity was affected upon mitochondrial and glycolysis block, while spine calcium influx was unaffected (Fig. 2E, S2D). These data suggest that the instant energy supply upon synaptic plasticity induction largely depends on mitochondrial oxidative phosphorylation. In contrast, glycolysis and mitochondrial oxidative phosphorylation provide sustained energy post-plasticity induction for at least 11 minutes. In the absence of the instant and sustained energy supply from mitochondria and glycolysis, synaptic plasticity is abolished.

As mitochondrial compartments are ∼30 μm long and support clustered plasticity in uninduced neighboring spines within the same dendrite^2,3^, we examined if the increase in spine ATP levels supported by mitochondria is restricted to the plasticity-induced spine or also made available to neighboring spines. Upon plasticity induction, spines adjacent to the plasticity-induced spine within 10 μm showed a statistically significant increase in ATP_spine_ compared to spines within 10-40 μm distance from the plasticity-induced spine (Fig. 2F). These data suggest that spine ATP increase is restricted to 10 μm of the plasticity-induced spine. This restricted spine ATP increase is not due to spine calcium spread from the plasticity-induced spine to the adjacent spines or direct stimulation of adjacent spines, as adjacent spines showed significantly reduced spine calcium influx compared to the plasticity-induced spine (Fig. S2E). These data suggested that the ∼10 μm restricted distribution of ATP_spine_ from the plasticity-induced spine can be achieved by a local distribution of ATP from the mitochondrial compartment at the base of the plasticity-induced spine.

### Synaptic plasticity induction drives local ATP_mito_ synthesis

To determine if mitochondria at the base of the plasticity-induced spine provide the local ATP, we measured mitochondrial ATP by targeting the ATP reporter to the mitochondrial matrix, fusing luciferase (luc) to four repeats of COX8 signal peptide^17^ (see Methods). To calibrate for reporter expression level, we fused luc with a pH-, thermo-, and photostable fluorescent protein mCherry2 (mCh)^18^, allowing ratiometric luminescence to fluorescence (L/F) readouts (see Methods). We refer to this ratiometric mitochondrial ATP reporter as Mito-ATP and the mitochondrial ATP measurements as ATP_mito_ (Fig. 3A). pH correction for ATP_mito_ was done as previously described by separate mitochondrial pH measurements using a pH-sensitive reporter targeted to the mitochondrial matrix, mito-pHluorin^17,19^ (Fig. S3A, B, see Methods).

**Figure 3.**
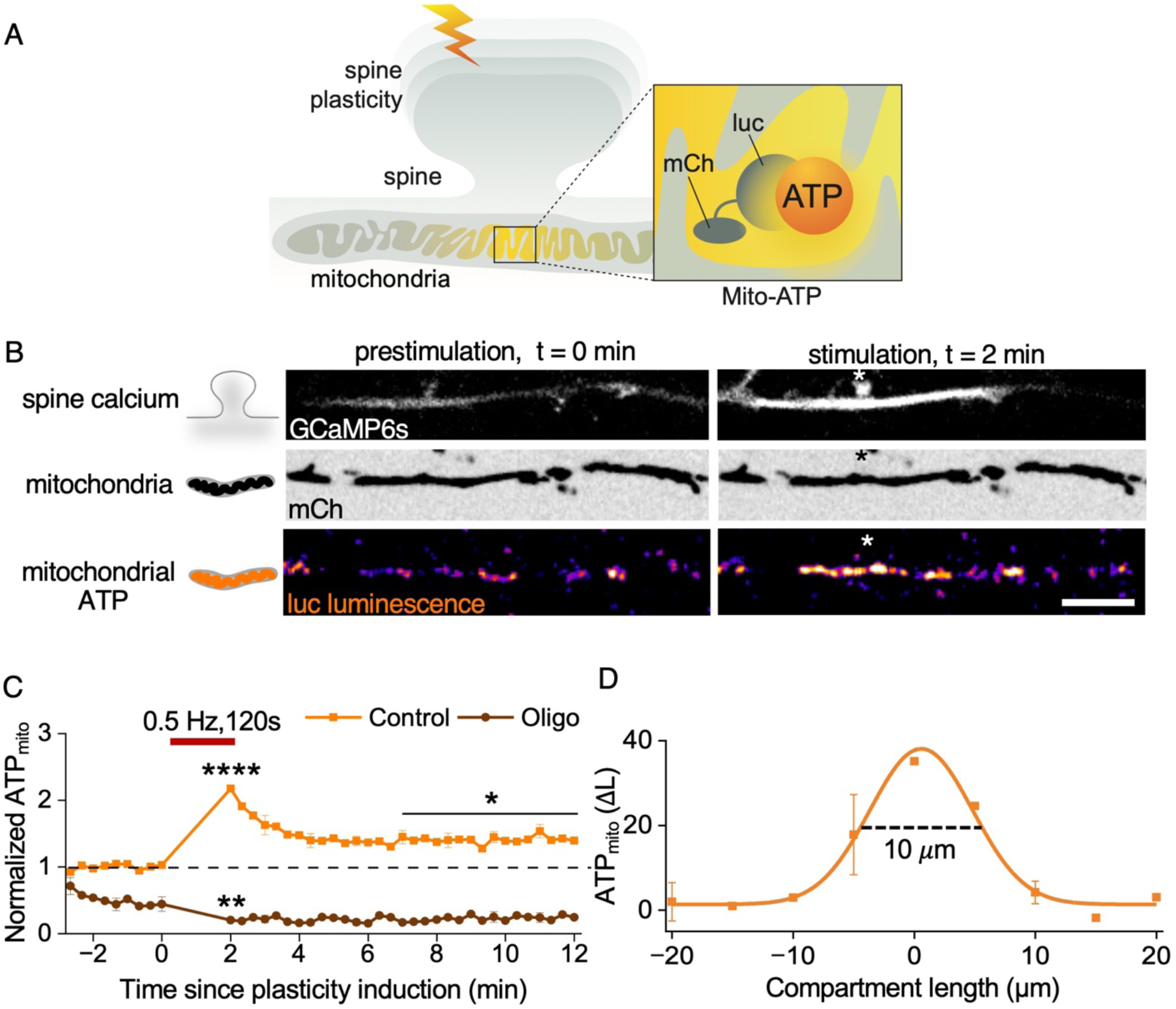
Synaptic plasticity induction drives instant, local, and sustained increase in ATP_mito_ A Illustration showing the design of the mitochondrial ATP sensor, Mito-ATP, for ATP_mito_ measurements following synaptic plasticity induction (lightning bolt). Luciferase (luc) is conjugated to the fluorescent protein mCherry2 (mCh) and targeted into the mitochondrial matrix (see Methods). **B** Representative images (of **C**, **D**) showing spine calcium influx in the plasticity-induced spine (white and black asterisk) measured using GCaMP6s fluorescence. In response to spine plasticity induction, there is a spatially restricted increase in mitochondrial luc luminescence at the base of the plasticity-induced spine without a change in the mCh fluorescence. Scale bar 10 μm. **C** The average time course of normalized ATP_mito_ (L/F) upon synaptic plasticity induction (red bar, 0.5 Hz, 120 s) shows an instant increase that declines and reaches a higher steady state (orange, Control). Upon ATP_mito_ synthesis inhibition (brown, Oligo), the instant and sustained increase in ATP_mito_ is lost. n in dendrites, animals: 14, 5 (Control), 7, 3 (Oligo). Paired Sample t test, p-values: <0.0001 (Control: Baseline vs. 2 min post-plasticity induction), 0.03219 (Control: Baseline vs. average of 6-11 min post-plasticity induction). Two sample t test, p-values: 0.00456 (2 min post-plasticity induction: Control vs. Oligo), 0.01959 (6-11 min post-plasticity induction: Control vs. Oligo). **D** Spatial profile of the change in mitochondrial luciferase luminescence (ΔL) along the dendrite shows a spatially restricted increase within 10 μm of the mitochondrial compartment at the base of the plasticity-induced spine. Negative distance values denote the direction of the cell body, whereas positive distance values denote the direction of the dendritic tip. n in dendrites, animals: 11, 3. See also Figure S3.

Expression of Mito-ATP in hippocampal neurons resulted in the efficient delivery of the reporter within the mitochondrial matrix, visualized by the mCh fluorescence and luc luminescence (Fig. 3A, B). Upon synaptic plasticity induction, we observed a rapid rise in ATP_mito_ in the first two minutes, which declined but remained sustained at a higher ATP_mito_ level for at least 12 minutes post-plasticity induction compared to baseline (Fig. 3B, C, Supp. Video 2), similar to the sustained rise in ATP_spine_ (Fig. 2C). This ATP_mito_ increase is generated by mitochondrial oxidative phosphorylation, as its inhibition (using Oligo) resulted in the depletion of ATP_mito_ (Fig. 3C). The ATP_mito_ increase upon synaptic plasticity induction was spatially restricted to a ∼10 μm compartment within the mitochondria at the base of the plasticity-induced spine (Fig. 3B, D, Supp. Video 2), in correspondence to the ∼10 μm spatial distribution of ATP_spine_ from the plasticity-induced spine (Fig. 2F). These results indicate that mitochondria respond to synaptic plasticity induction in time and space to generate instant, sustained, and local mitochondrial ATP.

### Synaptic input drives local mitochondrial calcium influx

In anticipation of the energy demands of synaptic plasticity and to drive local mitochondrial ATP synthesis, synapses should have a local signaling mechanism that links synaptic inputs to mitochondria. Mitochondrial calcium can activate enzymes of the Krebs cycle pathway, namely, pyruvate dehydrogenase, isocitrate dehydrogenase, and α-ketoglutarate dehydrogenase, to produce energy^20^. As there is substantial spine calcium influx during synaptic activity, we tested if calcium could be the local molecular signal that drives local mitochondrial ATP synthesis in response to synaptic input. While it is known that mitochondrial calcium entry drives mitochondrial ATP production^17,21^, it has not been examined in individual mitochondria near dendritic spines upon single spine stimulation.

To address this question, we measured mitochondrial calcium using a genetically encoded green calcium indicator, GCaMP6f, targeted to the mitochondrial matrix (MatrixGCaMP)^17^ (Fig. 4A). As control, we measured calcium outside mitochondria in its immediate vicinity, using GCaMP6f targeted to the outer mitochondrial membrane (OMMGCaMP)^17^ (Fig. 4B). Simultaneously, we monitored spine and dendritic calcium using a genetically encoded red calcium indicator, RCaMP1.07 targeted to the cytosol (RCaMP)^22^ (Fig. 4A, B). Spines were stimulated with 30 two-photon glutamate uncaging pulses at 0.3 Hz, which resulted in spine calcium influxes corresponding to each pulse. Interestingly, each spine calcium influx resulted in a temporally synchronous mitochondrial calcium influx (using MatrixGCaMP) (Fig. 4A, C, Supp. Video 3). The first pulse resulted in a large mitochondrial calcium signal sustained on successive pulses, in contrast to the spine calcium signal that declined to baseline at the end of each pulse (Fig. 4C, Supp. Video 3). On the other hand, calcium measured outside mitochondria in its immediate vicinity (using OMMGCaMP) showed a faster decay, resembling that of spine calcium response (Fig. 4B, C, Supp. Video 3). These data suggest that calcium follows distinct kinetics upon entering the mitochondrial matrix with slower clearance, and the difference in clearance kinetics between the mitochondrial matrix and outer mitochondrial membrane is not a property of the GCaMP6f calcium reporter.

**Figure 4.**
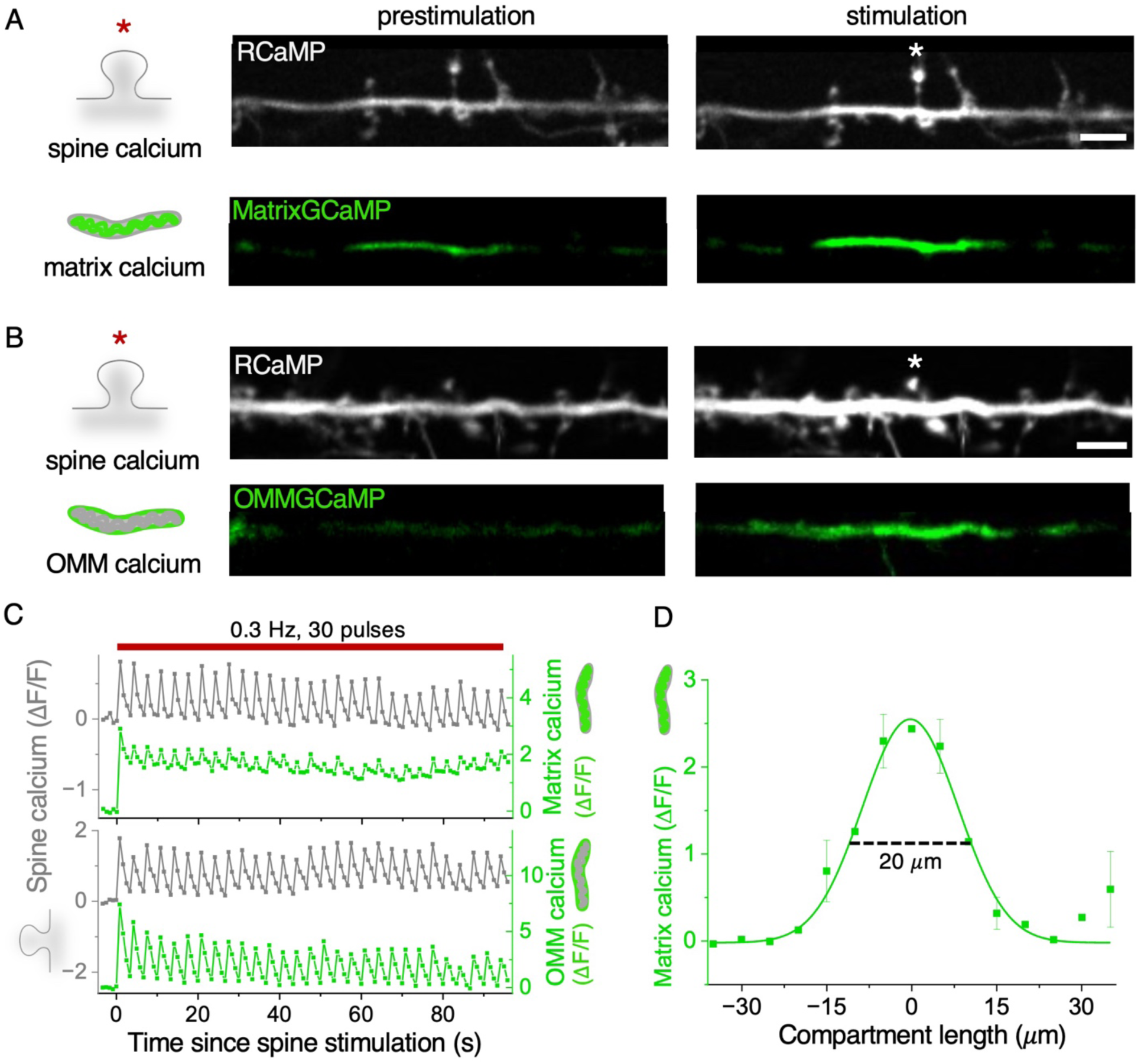
Spine stimulation drives instant, local, and sustained increase in mitochondrial calcium **A** Representative images (of **C**, **D**) showing an increase in spine calcium (white, RCaMP) upon single spine stimulation (white asterisk), followed by an instant and spatially restricted increase in mitochondrial matrix calcium (green, MatrixGCaMP) at the base of the stimulated spine. Scale bar 5 μm. **B** Representative images showing an increase in spine calcium (white, RCaMP) upon single spine stimulation (white asterisk), followed by a diffused increase in mitochondrial outer membrane calcium (green, OMMGCaMP) at the base of the stimulated spine. Scale bar 5 μm. **C** Representative time course (of **D**) of the spine (gray), mitochondrial matrix (green), and OMM calcium (green) responses (ΔF/F) during spine stimulation (red bar, 0.3 Hz 30 pulses) showing a large increase in mitochondrial matrix calcium that is sustained and responsive to each successive stimulation pulses, while the spine and OMM calcium are not sustained and decline to baseline at the end of each pulse. **D** Spatial profile of the change in mitochondrial matrix calcium (ΔF/F) along the dendrite shows a spatially restricted increase within 20 μm of the mitochondrial compartment at the base of the stimulated spine. Negative distance values denote the direction of the cell body, whereas positive distance values denote the direction of the dendritic tip. n in dendrites, animals: 13, 5. See also Figure S4.

Mitochondrial calcium influx upon spine stimulation is NMDAR-dependent, as blocking NMDAR (using APV) abolished spine and mitochondrial calcium response. (Fig. S4A, B). On the other hand, mitochondrial calcium influx upon spine stimulation is largely L-type calcium channel independent, as blocking the L-type calcium channel (using Nimodipine) showed a significant but small effect on the spine and mitochondrial calcium response (Fig. S4C, D). Mitochondrial calcium uptake was only observed when the dendritic calcium crossed a specific threshold, beyond which the amplitude of mitochondrial calcium uptake was positively correlated to dendritic calcium amplitude (Fig. S4E). Notably, when the calcium influx was restricted to only the spine head, by reducing the two-photon glutamate uncaging laser power, mitochondrial calcium uptake was not observed. These data indicate that the dendritic calcium must cross a specific threshold at the base of the stimulated spine to drive local mitochondrial calcium influx, as observed in axonal terminals^17^.

The temporal synchronicity in mitochondrial calcium uptake with spine and dendritic calcium suggests that calcium could be the local signaling molecule that links synaptic inputs’ energy demands to mitochondrial energy supply. Moreover, in correspondence to the spatially restricted mitochondrial ATP generation and spine ATP increase (Fig. 2B, F, 3B, D, Supp. Videos 1, 2), mitochondrial calcium uptake was also spatially restricted to ∼20 μm of the stimulated spine (Fig. 4A, D, Supp. Video 3). These data suggest that calcium could be the instant, local, and sustained signaling molecule that drives the instant, local, and sustained mitochondrial ATP synthesis and spine ATP increase in response to synaptic input.

### ER plays a regulatory, but not obligatory, role in mitochondrial calcium influx

To investigate if local mitochondrial calcium entry upon synaptic input drives mitochondrial ATP synthesis at the base of the plasticity-induced spine, we examined the mechanism of mitochondrial calcium entry. Local microdomains of high calcium concentration within ER are known to release calcium into mitochondria^23–26^. Disruption of ER-mitochondria tethering also affects mitochondrial calcium handling^2,27^. Therefore, we hypothesized that ER provides calcium to mitochondria upon spine stimulation. To test this hypothesis, we blocked the two main ER calcium regulatory pathways: (1) the Sarco/Endoplasmic Reticulum Calcium ATPase (SERCA) pump required for ER calcium uptake (using Thapsigargin or Cyclopiazonic acid (CPA))^17,28^; (2) Ryanodine receptors (RyR) and IP3 receptors (IP3R) required for ER calcium release (using Ryanodine and Xestospongin C (XestC), respectively)^28,29^ (Fig. 5A).

**Figure 5.**
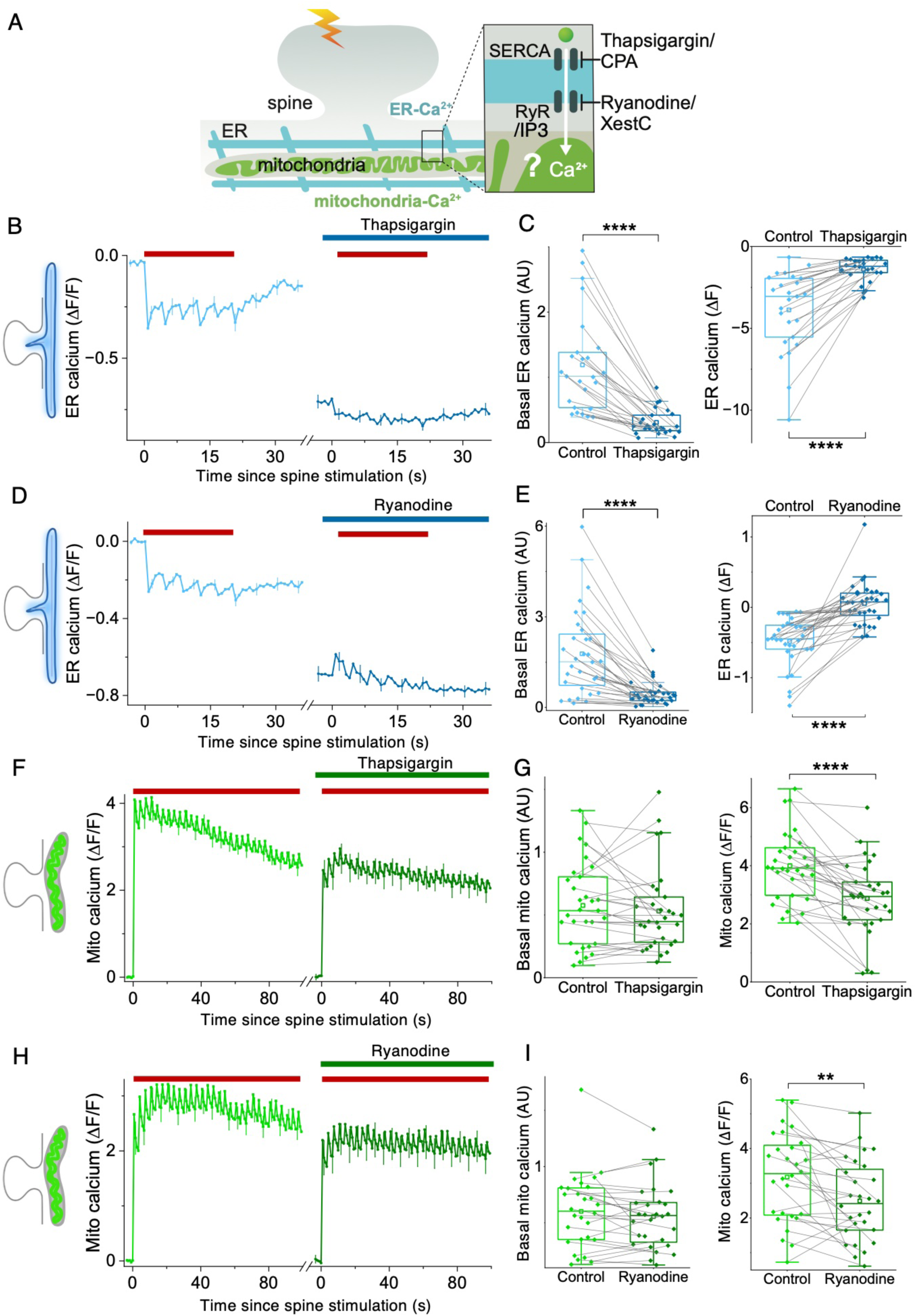
Local mitochondrial calcium entry upon spine stimulation is largely independent of ER **A** Illustration of ER (blue, ER-GCaMP210) and mitochondrial calcium (green, MatrixGCaMP) measurements upon spine stimulation (lightning bolt) following the inhibition of ER calcium influx (using Thapsigargin and CPA) and release (using Ryanodine and XestC). **B, D** The average time course of ER calcium response (ΔF/F) during spine stimulation (red bar, 0.25 Hz, 6 pulses) shows depletion of ER calcium stores upon Thapsigargin treatment and accumulation of ER calcium upon Ryanodine treatment. n in dendrites, animals: 25, 3 (Thapsigargin), 32, 3 (Ryanodine) **C, E** Basal ER calcium and ER calcium release (ΔF) upon spine stimulation is significantly decreased upon Thapsigargin and Ryanodine treatment. n in dendrites, animals: 25, 3 (Thapsigargin), 32, 3 (Ryanodine). Paired Sample Sign Test, p-value: <0.0001 (Basal ER calcium and ER calcium response: Control vs. Thapsigargin and Control vs. Ryanodine). **F, H** The average time course of mitochondrial calcium response (ΔF/F) during spine stimulation (red bar, 0.3 Hz, 30 pulses) shows a small decrease in mitochondrial calcium response upon Thapsigargin and Ryanodine treatments. n in dendrites, animals: 31, 3 (Thapsigargin), 28, 3 (Ryanodine). **G, I** Basal mitochondrial calcium is unaffected, and mitochondrial calcium response (ΔF/F) upon spine stimulation shows a small but significant decrease upon Thapsigargin and Ryanodine treatment. n in dendrites, animals: 31, 3 (Thapsigargin), 28, 3 (Ryanodine). Paired Sample t Test, p-values: 0.2812 (Basal mito calcium: Control vs. Thapsigargin,), <0.0001 (Mito calcium response: Control vs. Thapsigargin), 0.11229 (Basal mito calcium: Control vs. Ryanodine), 0.00146 (Mito calcium response: Control vs. Ryanodine). AU: arbitrary unit. See also Figure S5.

To confirm the efficient block of ER calcium uptake and release by these inhibitors, we measured ER calcium upon spine stimulation using a genetically encoded, ER-sensitive calcium indicator, GCaMP210, targeted to the ER lumen^30^. Interestingly, the regular spine stimulation protocol used to measure mitochondrial calcium influx (0.3 Hz, 30 pulses, 10 ms pulse duration) did not effectively release ER calcium^2^, indicating that mitochondrial calcium influx might be ER-independent. To further examine this observation and to confirm the effective inhibition of ER uptake and release, we used an uncaging protocol with 6 two-photon glutamate uncaging pulses at 0.25 Hz at 100 ms pulse duration, which resulted in synchronous ER calcium release for each uncaging pulse, as observed before^29,31^ (Fig. 5B, D, S5A, C, see Methods). Upon inhibition of ER calcium uptake (using Thapsigargin or CPA) or release (using Ryanodine) followed by spine stimulation, basal ER calcium content and release were effectively abolished, as in axonal terminals^30^, with some accumulation of ER calcium in response to each uncaging pulse in the presence of Ryanodine, as in dendrites^29^ (Fig. 5A-E, S5A, B). However, inhibition of IP3R (using XestC) did not affect ER calcium release (Fig. S5C, D).

Despite the complete depletion of ER calcium by blocking uptake or release, it only resulted in a small effect on mitochondrial calcium uptake and spine calcium response upon spine stimulation at 0.3 Hz, 30 pulses (Figure 5F-I, S5E-J), indicating that ER calcium does not largely influence mitochondrial calcium entry. These data suggest that ER serves a regulatory, but not obligatory, role in mitochondrial calcium entry upon spine stimulation (see Discussion).

### Local mitochondrial stabilization and calcium signaling enable local ATP_mito_ synthesis to fuel synaptic plasticity

As mitochondrial calcium uptake is largely ER-independent, we directly targeted the mitochondrial calcium uniporter (MCU)^17,32,33^ to examine the significance of local mitochondrial calcium entry in local mitochondrial ATP synthesis. Knocking down MCU reduces presynaptic mitochondrial calcium uptake upon whole-field neuronal stimulation^17^, but MCU’s role in dendritic mitochondrial calcium uptake is unknown. We perturbed MCU by using a previously characterized shRNA^17^ (Fig. 6A). MCU knockdown abolished mitochondrial calcium uptake upon spine stimulation but was accompanied by significantly reduced basal mitochondrial calcium (Fig. S5I, S6A, B).

**Figure 6.**
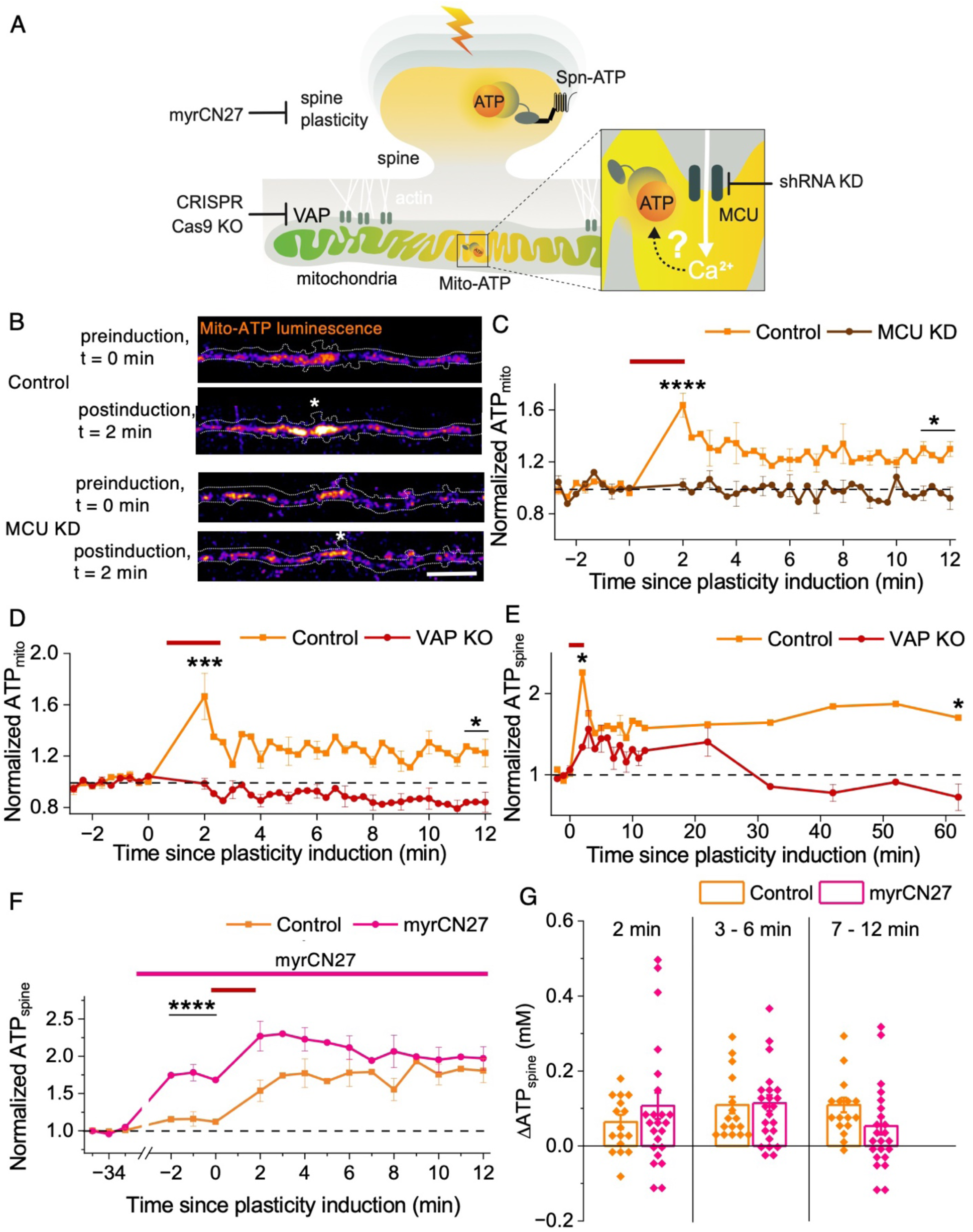
Local mitochondrial stabilization and calcium signaling drive local ATP_mito_ synthesis to fuel synaptic plasticity **A** Illustration of mitochondrial (yellow, Mito-ATP) and spine (yellow, Spn-ATP) ATP measurements upon synaptic plasticity induction (lightning bolt) following the inhibition of mitochondrial calcium uptake (using MCU shRNA) or mitochondrial stabilization (using VAP CRISPR Cas9 KO) or CaMKII-induced structural plasticity (using myrCN27). **B** Representative images (of **C**) showing Mito-ATP luminescence increase at the base of the plasticity-induced spine (white asterisk, dendrite trace in dotted white line) in the Control, but not in MCU KD neurons. Scale bar 10 μm. **C** The average time course of normalized ATP_mito_ upon synaptic plasticity induction (red bar, 0.5 Hz, 120 s) shows an instant increase in the Control (orange) that declines and reaches a higher steady state than baseline, but not in MCU KD (brown) neurons. n in dendrites, animals: 11, 6 (Control), 7, 3 (MCU KD). One-way ANOVA, Tukey test, p-values: <0.0001 (2 min post-plasticity induction: Control vs. MCU KD), 0.03372 (11-12 min post-plasticity induction: Control vs. MCU KD). **D** The average time course of normalized ATP_mito_ upon synaptic plasticity induction (red bar, 0.5 Hz, 120 s) shows an instant and sustained increase in the Control (orange, Cas9 Control) but not in VAP KO (red). n in dendrites, animals: 6, 3 (Control), 6, 5 (VAP KO). One-way ANOVA, Tukey test, p-values: 0.00029 (2 min post-plasticity induction: Control vs. VAP KO), 0.0418 (11-12 min post-plasticity induction: Control vs. VAP KO). **E** The average time course of normalized ATP_spine_ upon synaptic plasticity induction (red bar, 0.5 Hz, 120 s) shows an instant increase in the Control (orange, Cas9 Control) that declines and reaches a higher steady state than baseline at 62 min post-plasticity induction, but not in VAP KO (red). n in spines, animals: 10, 7 (Control), 8, 7 (VAP KO). One-way ANOVA, Tukey test, p-values: 0.04544 (2 min post-plasticity induction: Control vs. VAP KO), 0.02824 (62 min post-plasticity induction: Control vs VAP KO), 0.17906 (VAP KO: Baseline vs. 2 min post-plasticity induction). **F** The average time course of normalized ATP_spine_ shows an increase in baseline ATP_spine_ upon myrCN27 treatment (pink) and a similar increase as Control (orange) upon spine plasticity induction (red bar, 0.5 Hz, 120 s). n in spines, animals: 16, 3 (Control), 23, 7 (myrCN27). One-way ANOVA, Tukey test, p-values: < 0.0001 (Baseline −2 to 0 min: Control vs myrCN27). **G** The average change in ATP_spine_ (ΔATP_spine_) shows a similar increase upon myrCN27 treatment (pink) compared to the Control (orange) 2 min, 3-6 min, and 7-12 post-plasticity induction. n in spines, animals: 16, 3 (Control), 23, 7 (myrCN27). One-way ANOVA, Tukey test, p-values: 0.85314 (2 min post-plasticity induction: Control vs. myrCN27), 0.99999 (3-6 min post-plasticity induction: Control vs. myrCN27), 0.65015 (7-12 min post-plasticity induction: Control vs. myrCN27). See also Figure S6.

To determine whether the observed local ATP_mito_ synthesis (Fig. 3B-D, Supp. Video 2) is driven by local mitochondrial calcium entry (Fig. 4A, C, D, Supp. Video 3), we blocked mitochondrial calcium entry (using MCU KD) (Fig. S6A, B) and measured ATP_mito_ upon synaptic plasticity induction. The baseline ATP_mito_ was significantly increased in MCU KD neurons compared to the Control (Fig. S6C). Upon synaptic plasticity induction, in the absence of mitochondrial calcium entry (using MCU KD), the instant and sustained generation of mitochondrial ATP was abolished (Fig. 6B, C). The inability to generate local mitochondrial ATP synthesis was accompanied by deficits in synaptic plasticity formation and sustenance; however, spine calcium influx was also affected (Fig. S6D, E), indicating that the synaptic plasticity deficits observed in MCU KD spines could be due to the reduced spine calcium influx.

Therefore, we opted for another method to examine the significance of locally stable mitochondrial compartments^2,3^ in enabling the local calcium signal to drive ATP synthesis and support synaptic plasticity. We have shown that the Vesicle Associated Membrane Protein-Associated Protein (VAP), linked to the motor neuron disease Amyotrophic Lateral Sclerosis, plays a critical role in stabilizing dendritic mitochondria near synapses to support plasticity sustenance in stimulated and neighboring spines^2^. In the absence of VAP, mitochondrial calcium influx at the base of the plasticity-induced spine is significantly reduced without affecting spine calcium influx^2^. We, therefore, used VAP as a molecular handle to destabilize mitochondria and local mitochondrial calcium influx and examine mitochondria’s ability to respond to synaptic calcium, drive local mitochondrial ATP synthesis, and sustain the heightened spine ATP. We depleted VAP using CRISPR-Cas9, which effectively destabilizes mitochondrial compartments in dendrites^2^, and measured ATP_mito_ and ATP_spine_ levels upon synaptic plasticity induction. The baseline ATP_mito_ significantly increased in VAP KO compared to the Control, while ATP_spine_ showed an increasing trend (Fig. S6F, G). Upon synaptic plasticity induction, unstable mitochondria in VAP-depleted neurons could not generate spatially localized ATP_mito_ at the base of the plasticity-induced spine compared to stable mitochondria in Control neurons (Fig. 6D). Similarly, while the Control spines exhibited increased and sustained ATP_spine_ for up to an hour following synaptic plasticity induction, spines in VAP-depleted neurons with unstable mitochondria did not exhibit heightened and sustained ATP_spine_ (Fig. 6E). While VAP-depleted neurons showed a slight and statistically insignificant increase in ATP_spine_ 2 minutes post-plasticity induction, potentially supported by glycolysis (Fig. 2C, D), it was not sustained for an hour as in Control neurons (Fig. 6E). These data indicate that local mitochondrial stabilization is critical to enabling the local mitochondrial calcium signaling to drive sustained ATP_mito_ synthesis and heightened ATP_spine_ to support synaptic plasticity.

Next, we examined whether this local calcium signaling to drive spine-specific mitochondrial ATP synthesis is driven by CaMKII-dependent structural plasticity and its associated energy demands. CaMKII, the calcium-calmodulin (CaM)-dependent protein kinase II protein, plays a central role in synaptic plasticity and learning^34^. It is a serine/threonine protein kinase with 12 subunits; each subunit consumes ATP for its autophosphorylation and self-activation upon synaptic plasticity induction^35^. Besides ATP consumption for CaMKII kinase activity, CaMKII activation drives many energy-consuming downstream signaling processes, including actin polymerization and AMPAR trafficking and insertion within dendritic spines essential for synaptic plasticity formation and maintenance^36^. We blocked CaMKII and its downstream energy-demanding processes using the CaMKII inhibitor, myristoylated CN27 (myrCN27)^37^. The effective inhibition of CaMKII inhibition was confirmed by the lack of spine structural plasticity upon myrCN27 treatment (Fig. S6H). We expected that upon CaMKII inhibition, synapses would have negligible energy demand due to the absence of spine structural plasticity, which, therefore, will abort the local calcium signaling and the subsequent local mitochondrial ATP synthesis. Upon CaMKII inhibition, baseline ATP_spine_ significantly increased compared to the Control, suggesting that CaMKII activity and its downstream processes consume energy even in the presence of TTX (Fig. 6F, see Methods). Interestingly, upon synaptic plasticity induction, CaMKII-inhibited spines showed a similar increase in ATP_spine_ 2-6 min post-plasticity induction compared to the Control, with a decreasing trend 7-12 min post-plasticity induction compared to the Control (Fig. 6F, G). These results indicate that the initial burst of spine ATP increase, supported by mitochondrial ATP synthesis (Fig. 2C, D, 3B, C, 6B-E), is independent of CaMKII and its associated energy demands of spine structural plasticity. It also suggests that the initial burst of spine ATP increase upon plasticity induction might be anticipatory in nature.

In summary, these data indicate that the synapse-driven local mitochondrial calcium entry enabled by mitochondrial stabilization is critical to drive the instant and sustained local mitochondrial ATP production and the sustained increase in spine ATP to support synaptic plasticity (Fig. 7).

**Figure 7.**
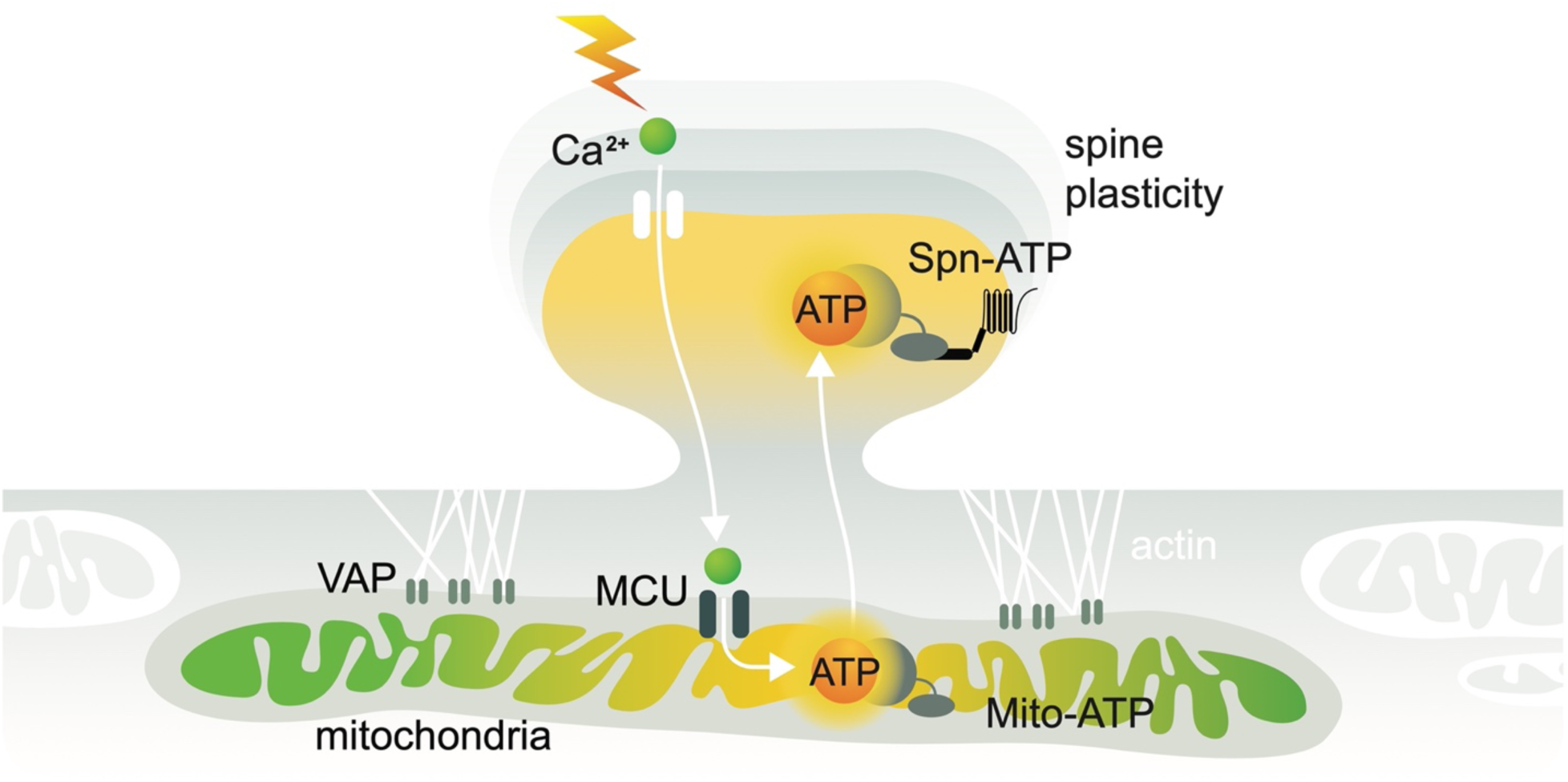
Synapses drive local mitochondrial ATP synthesis to fuel plasticity Illustration showing the significance of synaptic calcium signal input (green sphere) driving local mitochondrial calcium signaling (green sphere, green cristae) enabled by local mitochondrial stabilization (VAP, actin tethers) resulting in spatially restricted mitochondrial ATP production (yellow cristae) and its distribution to plasticity-induced spines (yellow spines) measured using mitochondria-(Mito-ATP) and spine-targeted (Spn-ATP) ATP reporters, respectively.

## Discussion

Even brief disruptions in energy supply can lead to cognitive deficits and brain disorders^1,4,5^. Remarkably, the energy producers, mitochondria, are prime suspects in their inability to generate adequate energy for synaptic demands in many neurological disorders such as Alzheimer’s, Parkinson’s, and Amyotrophic Lateral Sclerosis^1,4,5^. Using state-of-the-art ATP reporters and imaging methods to visualize and quantify ATP at single-spine and -mitochondrial resolution upon single-spine stimulation and plasticity induction using two-photon glutamate uncaging, we probe various fundamental mechanisms driving energy provision to synapses. We demonstrate the following: (1) steady-state spine ATP levels correspond to 10^5^ molecules in a typical dendritic spine and are tightly linked to the activity state of neurons and spines, supported by glycolysis and mitochondrial oxidative phosphorylation; (2) upon synaptic plasticity induction, spines ramp-up their ATP levels 1.5 fold within 10 μm from the plasticity-induced spine; (3) mitochondria generate the instant and sustained increase in spine ATP within ∼10 μm spatial compartments at the base of the plasticity-induced spine; (4) local mitochondrial ATP generation at the base of the plasticity-induced spine is driven by local mitochondrial calcium entry, independent of the endoplasmic reticulum; (5) mitochondrial stabilization near synapses is essential for the local calcium signaling and the instant, sustained mitochondrial ATP generation and heightened spine ATP to support plasticity; (6) the initial burst of spine ATP supported by mitochondrial ATP synthesis is not driven by energy demand and might be anticipatory.

Using a modified version of a previous presynaptic ATP reporter^6^ and a custom-built microscope, we have quantified the absolute steady-state ATP levels in individual spines. We find that a typical hippocampal pyramidal neuron’s steady-state ATP_spine_ is ∼0.4 mM (Fig. 1C, S1C), three times lower than a typical hippocampal pyramidal neuron’s presynaptic terminal ATP_presyn_ ∼1.4 mM^6^. This difference might stem from the innate differences in the energy-consuming processes and the energy-supplying mechanisms between dendritic spines and axonal terminals, such as: (1) dendritic spines are innately tuned to receive and store information, whereas axonal terminals are tuned to transmit information; therefore, the molecular machinery in each of these compartments must have different energy needs and timescales at which they function; (2) dendritic spines are highly reliant on mitochondria for their most energy-demanding process such as local protein synthesis, synaptic plasticity induction and maintenance, with minimal input from glycolysis^3,38^ (Fig. 2C, D, 6E); meanwhile, axonal terminals are reliant on both mitochondria and glycolysis during their most energy demanding process such as complete recycling of their total synaptic vesicle pool^6^; consistent with this notion, mitochondria in dendrites are much longer ∼30 μm and stable compared to mitochondria in axons ∼1 μm and motile^3,39,40^, indicating the different adaptations of the energy supplies based on the extent and frequency of energy burden in these two neuronal compartments; (3) upon plasticity induction, dendritic spine ATP levels ramp up and are sustained at ∼1.5 fold higher than baseline (Fig. 2C, 6E, F), perhaps to compensate for their low starting steady-state ATP levels; whereas upon synaptic stimulation, axonal terminal ATP levels remain unchanged^6^ or show a slight ∼10-20% decrease^15,16^, perhaps due to their high steady-state ATP levels, revealing different sensitivities to energy needs and rates of energy synthesis between the two compartments; (4) the steady-state spine ATP is highly heterogeneous across neurons with a coefficient of variation (CV) of 67% (Fig. S1C), which is higher than in axonal terminals measured across neurons with a CV of 29%^6^, suggesting that the spines’ activity history might determine their steady-state ATP levels. However, spine size, which could be used as one of the correlates of spine activity history, did not show a strong correlation with steady-state spine ATP levels (Fig. S1F). We also observed that spine ATP decreased with increasing distance from the soma (Fig. S1E). This result suggests that low steady-state spine ATP levels may suppress specific ATP-dependent mechanisms within distal spines, which could be activated as spine ATP levels increase upon plasticity induction. These data further emphasize the need for local energy supplies in distal spines to drive specific ATP-dependent mechanisms and should be investigated in the future.

The state-of-the-art ATP imaging methods developed here have allowed us to unravel the intricate differences in the energetics of dendritic spines compared to axonal terminals, which will be extremely valuable in building theoretical synaptic and dendritic computation models. The mechanisms underlying these differences, their molecular determinants, and the role of other energy sources besides mitochondria and glycolysis, such as glial-lactate shuttle^41^, lipid droplets^42^, and glycogen granules^43^, should be further elucidated with the help of these and recent methods to measure ATP^15^.

We previously showed that dendritic mitochondria form 30 μm long stable compartments to sustain synaptic plasticity for up to an hour and determine the spatial extent of neighboring spines supported within the dendrite^2^. However, it has been challenging to visualize the energy currency, ATP, their synthesis rate within mitochondria, and their distribution to dendritic spines. With the newly developed mitochondria- and spine-targeted ATP reporters, we have directly measured ATP synthesis and its distribution within 10 μm of the plasticity-induced spine (Fig. 2B-D, F, 3B-D, Supp. Videos 1, 2). These results corroborate our previous findings that local impairment of a mitochondrial compartment affects plasticity-induced protein synthesis only in spines served by the defective mitochondrial compartment^3^. In contrast, the plasticity-induced protein synthesis in neighboring spines with healthy mitochondrial compartments remains unaffected^3^. In the future, the mechanisms that spatially restrict the locally generated mitochondrial ATP within 10 μm of the plasticity-induced spine need to be examined. Following synaptic plasticity induction, we see a decline in the instantly generated mitochondrial ATP before it reaches a higher steady state than baseline (Fig. 3C), indicating a sustained release of ATP from mitochondria to the cytosol through the ADP/ATP carriers on the mitochondrial inner membrane^44,45^. Super-resolution microscopy would be critical to reveal whether a spatial restriction or recruitment of the ADP/ATP carriers on the inner mitochondrial membrane at the base of the plasticity-induced spine drives the spatially restricted distribution of ATP to plasticity-induced spines within 10 μm. Super-resolution methods could also be used to examine if the mitochondrial calcium gating machinery, the MCU complex, is spatially restricted or recruited to the base of the plasticity-induced spine to enable local mitochondrial calcium entry and ATP synthesis in anticipation of energy demands. These spatially restricted mechanisms further highlight the importance of local energy sources in supporting synapses, as mere ATP diffusion cannot meet the instant and sustained ATP needs at the optimal ATP concentrations required to drive most synaptic molecular processes^6,46^.

The sustenance of synaptic plasticity and the spread of plasticity to adjacent spines within 10 μm is supported by the sharing of locally available and newly synthesized proteins, called ‘plasticity-related proteins’, between spines^47–49^. Our findings showing the dependence of plasticity-induced spines on sustained ATP and its distribution within 10 μm indicate that dendritic spine function is likely limited by ATP availability. It, therefore, opens the possibility that ATP could be used as a plasticity tag or an obligatory fuel to drive the function of some plasticity-related proteins, most of which are ATP-consuming enzymes (CaMKIIa^34^, actin^50^, cofilin^51^). Future studies should, therefore, investigate if ATP-rich synapses could be used as markers for synaptic potentiation following a learning paradigm *in vivo*.

ER is the primary calcium storage organelle, and ER-mitochondria contacts (ERMC) are essential for mitochondrial calcium dynamics in neurons^2,27,52^. However, the standard spine stimulation protocol required to drive mitochondrial calcium entry, mitochondrial ATP synthesis, spine ATP synthesis, and spine plasticity failed to induce substantial ER calcium release^2^ (Fig. 2B-F, 3B-D, 4A, C, D, S4A-D). While upon using a modified spine stimulation protocol, ER calcium release was observed at the base of the stimulated spine, neither blocking ER calcium intake (using Thapsigargin or CPA) nor release (using Ryanodine) substantially affected mitochondrial calcium uptake (Fig. 5F-I, S5E-I). These results corroborate the previous finding that mitochondrial calcium uptake is insensitive to blocking ER calcium release within axonal terminals (using CPA)^17^. On the contrary, disruption of ER-mitochondria tethering dramatically affects mitochondrial calcium uptake^27,53^. Recent work also reveals that ER to mitochondrial calcium transfer only occurs when enough ER calcium is released^31^. These observations suggest that while ER-mitochondria tethering might play a role in mitochondrial calcium uptake, ER-mitochondria tethering is critical for stabilizing mitochondria at the base of spines to enable local mitochondrial calcium entry in the absence of ER or sufficient ER calcium release. This notion is further supported by the electron microscopy observation that only a small subset of dendritic spines have ER in hippocampal pyramidal neurons^54,55^, indicating that ER is not compulsory to transmit spine calcium signal to local mitochondria to drive ATP production and synaptic plasticity. Notably, to effectively communicate the spine calcium signal to local mitochondria, the spine calcium needs to enter the dendritic shaft at the base of the stimulated spine and cross a specific threshold to enable mitochondrial calcium influx (Fig. S4E).

In the motor neuron disease ALS, in the absence of stable mitochondria due to the defective ER and mitochondria-actin tethering protein VAP, the spines lose their ability to sustain plasticity^2^. Our new ATP imaging methods reveal the spine and mitochondrial energetics of the ALS-linked, VAP-deficient neurons. We find that the sustained reliance of spines on sustained mitochondrial ATP is essential for synaptic plasticity, the absence of which results in energy deficits potentially leading to ALS, its associated learning deficits, and other neurological disorders. The sustained increase in mitochondrial and spine ATP levels might also play an essential role in the sustained mitochondrial transcription and the subsequent protein synthesis required to improve mitochondrial function in aged mice^56^. These advanced ATP imaging methods to quantify steady-state ATP levels and their dynamics at single-spine and mitochondrial resolution allow new studies on energy homeostasis in healthy aging^57^ and identify new targets and therapeutics for devastating brain disorders.

## Acknowledgments

We thank H. Bito for the RCaMP1.07 plasmid; C. Hanus for the PSD95-mCherry plasmid; J. de Juan-Sanz for Homer2-mOrange2-luc plasmid; G. Ashrafi for mito-pHluorin and MCU shRNA plasmids; T. Ryan for MatrixGCaMP, OMMGCaMP, and ERGCaMP plasmids; D. Kim for GCaMP6s plasmid; and R. Yasuda for mEGFP plasmid. We are also thankful to J. Zur for mitochondrial and ER calcium image analysis scripts; S. Perez, D. Bhuyan, S. Yang, and G. Sathyanarayan for preparing cultured hippocampal neurons and technical assistance; L. Yan at the MPFI Imaging Center Core for assistance with custom-building the microscope; J. Yu at the MPFI Molecular Virology Core for assistance with molecular cloning; and R. Yasuda, H. Inagaki, and all members of the Rangaraju Lab for critical input on the manuscript. IG is funded by the Carl Angus Desantis Foundation; VR is supported by the Max Planck Society, Louis D. Srybnik Foundation, F.O.R.E Foundation, Chan Zuckerberg Initiative DAF an advised fund of the Silicon Valley Community Foundation grant numbers 2023-331775 and 2024-349543, and the NIH Director’s New Innovator Award (DP2 MH140148).

## Author Contributions

VR, IG, and RF designed experiments. Experiments were carried out by IG, RF, and OB. Data were analyzed by IG, RF, OB, and MS. VR, RF, and IG wrote the manuscript, and all authors edited the manuscript.

## Materials and Methods

### Animals

All experiments were performed according to the Max Planck Florida Institute for Neuroscience IACUC regulations (protocol number 22-005).

### Plasmid constructs

Mito-ATP reporter was developed by replacing the GCaMP6f in MatrixGCaMP^17^ with mCherry2 and luciferase from Syn-ATP^6^. RCaMP1.07 plasmid was obtained from the Bito Lab^22^, PSD95-mCherry plasmid from Hanus C^58,59^, Homer2-mOrange2-luciferase plasmid from the de Juan-Sanz Lab, mito-pHluorin plasmid and MCU shRNA plasmids from the Ashrafi Lab^17^, and mEGFP plasmid from the Yasuda Lab. The Homer2-EGFP-luciferase plasmid was made by replacing mOrange2 with EGFP in the Homer2-mOrange2-luciferase plasmid. MatrixGCaMP (GCaMP6f targeted to four repeats of COX8 signal peptide of the mitochondrial matrix)^17^, OMMGCaMP (GCaMP6f targeted to TOM20 signal peptide on the outer mitochondrial membrane)^17^, ERGCaMP (GCaMP6-210 targeted to N-terminal Calreticulin signal peptide)^30^, and GCaMP6s^60^ plasmids were purchased from Addgene (127870, 127874, 86919, 40753 respectively).

### Cell culture preparation and transfection

Unless specified otherwise, all reagents were purchased from Sigma, and all stock solutions were stored at −20 °C. Sprague-Dawley rats (postnatal P0) were obtained from the in-house animal core facility of the Max Planck Florida Institute for Neuroscience. Hippocampal regions were dissected in ACSF containing (in mM) 124 NaCl, 5 KCl, 1.3 MgSO_4_:7H2O, 1.25 NaH_2_PO_4_:H_2_O, 2 CaCl_2_, 26 NaHCO_3_, and 11 Glucose (stored at 4 °C) and stored in hibernate E buffer (BrainBits LLC, stored at 4°C). Dissected hippocampi were dissociated using the Papain Dissociation System (Worthington Biochemical Corporation, stored at 4 °C) with a modified manufacturer’s protocol. Briefly, hippocampi were digested in papain solution (20 units of papain per ml in 1 mM L-cysteine with 0.5 mM EDTA) supplemented with DNase I (final concentration 95 units per ml) and placed in a shaking heat block for 30 min at 37 °C, 900 rpm. Digested tissue was triturated and set for 5 min, and the supernatant devoid of tissue chunks was collected. The supernatant was centrifuged at 300 rcf for 5 min, and the pellet was resuspended in resuspension buffer (1 mg of ovomucoid inhibitor, 1 mg of albumin, and 95 units of DNase I per ml in EBSS). The cells were forced to pass through a discontinuous density gradient formed by the resuspension buffer and the Ovomucoid protease inhibitor (10 mg per ml) with bovine serum albumin (10 mg per ml) by centrifuging at 600 rpm for 6 min. The final cell pellet devoid of membrane fragments was resuspended in Neurobasal-A medium (Gibco, stored at 4°C) supplemented with Glutamax (Gibco, stored at −20 °C) and B27 supplement (Gibco, stored at −20°C). Cells were plated on poly-D-lysine coated coverslips mounted on MatTek dishes at a density of 70000-80000 cells/cm^2^. Cultures were maintained at 37 °C and 5% CO_2_ by feeding them with the same medium every 3-4 days until transfection. Transfections were performed 12-18 days after plating by magnetofection using Combimag (OZ biosciences, stored at 4°C) and Lipofectamine 2000 (Invitrogen, stored at 4°C) according to manufacturer’s instructions.

### Imaging and optical measurements

Live cell imaging was conducted between 13-21 days after plating. All experiments were performed at 37 °C in modified E4 imaging buffer containing (in mM) 120 NaCl, 3 KCl, 10 HEPES (buffered to pH 7.4), 4 CaCl_2_, and 10 Glucose, unless specified otherwise.

Imaging was performed using a custom-built inverted spinning disk confocal microscope (3i imaging systems; model CSU-W1) with two cameras: an Andor iXon Life 888 electron-multiplying charge-coupled device (EMCCD) camera for confocal fluorescence imaging and an Andor iXon3 888 camera for luminescence imaging. The Andor iXon3 888 EMCCD camera was selected for ultralow dark noise, further reduced by cooling to −100 °C. The speed of the Andor iXon3 888 camera used for luminescence measurements was 1 MHz with 1.00 gain and 1000 intensification. Image acquisition was controlled by SlideBook 6 software. Images were acquired with a Plan-Apochromat 63x/1.4 NA. Oil objective, M27 with DIC III prism, using a CSU-W1 Dichroic for 488/561 nm excitation with Quad emitter and individual emitters, at laser powers 50-100 ms exposure, 2 mW (488 nm) and 50-100 ms exposure, 5.5 mW (561 nm) for confocal fluorescence, and a 720 nm multiphoton short-pass emission filter for luminescence. During imaging, the temperature was maintained at 37 °C using an Okolab stage top incubator with temperature control. The temperature was maintained at 32 °C for spine structural plasticity experiments for MCU KD and related Control neurons.

### ATP_spine_ imaging and analysis

Neurons were transfected with Spn-ATP and GCaMP6s, along with the MCU shRNA, VAP sgRNA, or Cas9 control, when specified. Transfected neurons with mature spines were identified by a change in GCaMP6s fluorescence corresponding to calcium transients. In addition, the mOrg signal from Spn-ATP was used to examine its expression and identify spines for imaging.

Before glutamate uncaging and imaging, the imaging buffer was replaced with 2 mM luciferin (Promega) for the bioluminescence reaction to detect ATP in the spines, 1 μM TTX (Citrate salt, 2 mM stock made in water), 50 μM Forskolin (Tocris Bioscience, 25 mM stock made in DMSO), and 2 mM 4-Methoxy-7-nitroindolinyl-caged-L-glutamate (MNI caged glutamate) (Tocris Bioscience, 100 mM stock made in E4 buffer) in modified E4 buffer lacking Mg^2+^ (see above). For pharmacological inhibition of CaMKII, myrCN27 (1 µm)^61^ was added to the imaging buffer and incubated for 30 mins. Sample images of the selected neurons were acquired simultaneously in the 561 nm (mOrg) and 488 nm (GCaMP6s) channels at exposure times of 50 ms each, and bioluminescence was acquired over 60 s. The neuron with the best mOrg fluorescence was selected for further experiments. Dendritic branches 50-100 µm away from the soma, with mature mushroom-shaped spines showing reliable baseline luminescence, were chosen for two-photon glutamate uncaging. Glutamate uncaging was performed using a multiphoton-laser 720 nm (Chameleon, Coherent) and a Pockels cell (Conoptics) to control the uncaging pulses. To test a spine’s response to an uncaging pulse, an uncaging spot (2 μm^2^) close to a spine head was selected, and one to two uncaging pulses at 10 ms pulse duration per pixel and 7 mW back aperture power were given and checked for spine-specific calcium transients. For spine plasticity induction, an uncaging protocol of 60 uncaging pulses at 0.5 Hz with 10 ms pulse duration per pixel at 6 mW back aperture power of the 720 nm laser was used. An automated program was used for imaging the baseline (3 min), two-photon glutamate uncaging to induce synaptic plasticity induction (2 min), and post-plasticity induction (11 min) of the spine of interest with simultaneous acquisitions in 488 (GCaMP6s), 561 (mOrg) and luminescence (luciferase bioluminescence) channels using the same exposure and acquisition times as mentioned above, using the 6D feature of the Slidebook 6 software.

Images were analyzed using a customized ImageJ macro script to measure spine ATP at each time point. All time frames were concatenated into separate fluorescence and luminescence image stacks and corrected for XY drifts using the StackReg plugin. 5 μm diameter regions of interest (ROIs) were manually drawn to identify the stimulated and adjacent spines. Within these ROIs, the spines were segmented from the background into 2 μm diameter ROIs based on the pixels showing fluorescence or luminescence intensity. These ROIs were centered at the XY centroid of the spines to track the spines in case of any physical movement of the spines or dendrites during the time series. Background subtracted luminescence intensity (L) from the spine of interest was divided by the corresponding background subtracted mOrg fluorescence intensity (F) to determine the L/F ratio of the spine of interest at each time point as:

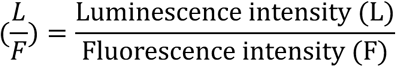

For measuring the spatial distribution of ATP, the above ImageJ macro script was used to concatenate images from each time points of baseline and plasticity induction (1 min), corrected for any drift in the XY plane using the StackReg Plugin, and ROIs were drawn around the spines within 40 µm from either side of the plasticity-induced spine as described above. Background subtracted L and F values and their corresponding L/F ratios were obtained as described above. The change in L/F (ΔL/F) was calculated by subtracting the L/F at the 1 min post-plasticity induction time point from the corresponding average baseline L/F of the same spine of interest.

### Spn-ATP calibration

Steady-state luminescence and fluorescence images were captured from the Spn-ATP-transfected neurons to record the baseline L/F ratio in live neurons in the modified E4 imaging buffer lacking Mg^2+^ (pH 7.4). For permeabilization, the imaging buffer was replaced with 5-10 U/mL Streptolysin-O (Millipore Sigma) prepared in PIPES buffer^6^ containing (in mM): 20 PIPES (buffered to pH 7.0 at 37 °C), 139 KCl, 0.91 EGTA, 0.186 CaCl_2_, and 5.22 MgCl_2_, for 2 min. The neurons were washed twice in PIPES buffer (pH 7.0) with 2 min incubation per wash to wash off intracellular ATP. Luminescence and fluorescence images were captured from the permeabilized neurons to record the L/F ratio at 0 mM ATP. Following permeabilization and wash, the luminescence intensity was negligible, indicating a complete wash-out of intracellular ATP. In contrast, the fluorescence intensity remained unchanged, confirming that the reporter was not washed out. For Spn-ATP calibration, permeabilized neurons were incubated in known ATP concentrations ranging from 0.1 to 5 mM prepared in PIPES buffer (pH 7.0) containing 2 mM luciferin at 37 °C. The obtained ATP titration curve of the L/F ratio vs. ATP concentrations was fit to the Michaelis-Menten equation below to determine the Michaelis-Menten constant, K_m_ of spine ATP, where [S] is the known ATP concentration, V is the L/F corresponding to the known ATP concentration, and V_max_ is the (L/F)_max_.

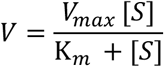

We performed these Spn-ATP calibration experiments once at the beginning of this project (Calibration 1: 2021-2023) and repeated it towards the end of this project (Calibration 2: 2024-2025) due to changes in the back-aperture laser power of the 561 nm laser in 2024, that affected mOrg fluorescence intensity. Consequently, Calibration 2 showed a slightly elevated L/F ratio due to decreased mOrg fluorescence, resulting in an elevated V_max_. Therefore, the mean spine ATP obtained from the two calibrations were comparable. Spines with L/F ratios above 5 times the standard deviation of the mean (8 spines out of a total of 395 spines in Calibration 1, 0 spines out of a total of 156 spines in Calibration 2, in Fig. 1B, C) were eliminated from further analysis as they could not be converted to reliable spine ATP concentrations.

### Estimates of ATP consumption by Spn-ATP

We calibrated our EMCCD to provide a conversion between luminescence intensity and the number of photons per unit luminescence intensity (15 photons/unit luminescence intensity; signal-to-noise ratio 10.5). Our luminescence measurements using Spn-ATP correspond to a total photon flux integrated over the area of a spine of 4320 photons/min. Considering 5% loss in photon collection efficiency due to the optical path and the published quantum yield of luciferase 0.88^62^, we are consuming 5168 ATPs/min/spine (assuming spine volume as 0.5 μm^3^), which is a small fraction, 4.2%, of the steady-state ATP levels (10^5^ ATPs/spine) and is 0.1% of the ATP production rate 6X10^6^ ATPs/min/spine^12^.

### pH correction for ATP_spine_ measurements

Cytosolic pH (pH_cytosol_) was determined using the pH-sensitive mOrg fluorescence at each time point or each experimental condition (DGlu+Pyr, Oligo, myrCN27, Cas9 Control, and VAP KO), with respect to the known baseline cytosolic pH of 6.9^6^ using a modified Henderson-Hasselbach equation:

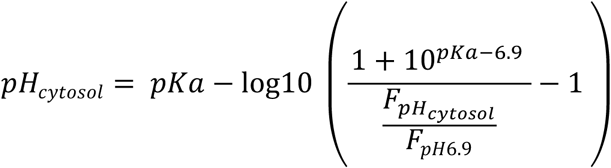

where 6.9 is the baseline cytosolic pH, pKa is 6.5 for mOrg, F_pH_ is the mOrg fluorescence intensity at each calculated pH, and F_pH6.9_ is the mOrg fluorescence intensity at baseline.

Since the ATP calibration of Spn-ATP was done at pH 7, the baseline fluorescence intensity was corrected from pH 6.9 to pH 7 for each experimental condition as follows:

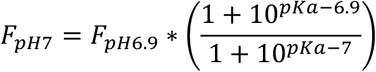

where 6.9 is the baseline cytosolic pH, pKa is 6.5 for mOrg, and *F*_pH_ is the mOrg fluorescence intensity at baseline pH 6.9.

F_pH7_ was then used to normalize the expression level of Spn-ATP at all time points following plasticity induction, and the L/F ratios were calculated as follows:

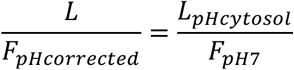

where *L_pHcytosol_* is the luminescence intensity at each observed pH.

The above L/F_pHcorrected_ ratios were converted to [ATP] knowing the impact of pH on the K_m_ of ATP^6^ using a modified Michaelis-Menten equation as:

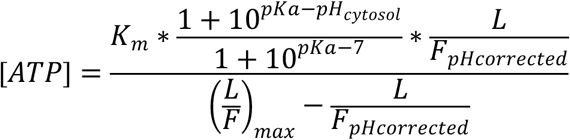

where the K_m_ and (L/F)_max_ or V_max_ are the Michaelis-Menten parameters obtained from the ATP titration curve measured at pH 7.0, and pK_a_ is pK_a_ of luciferase, 7.03. The effect of errors in pH measurement was not propagated to the standard error of the mean of [ATP].

### Measurement of Spn-ATP reporter movement or trafficking

To investigate the effect of spine plasticity induction on the movement or trafficking of Spn-ATP, we used a modified spine ATP reporter Homer2-EGFP-Luc, in which the mOrg of Spn-ATP was replaced with the pH-insensitive fluorescent protein EGFP. Neurons were co-transfected with Homer2-EGFP-Luc and RCaMP1.07 and imaged using the same glutamate uncaging protocol described for Spn-ATP above. We measured the EGFP fluorescence intensity, indicating the change in Homer2-EGFP-Luc levels, following plasticity induction in the Control, DGlu+Pyr, and Oligo-treated neurons as above. There was a decline in EGFP fluorescence intensity upon plasticity induction, which then recovered to baseline in all treatment conditions, indicating that the decline in the mOrg fluorescence intensity (F) of Spn-ATP upon plasticity induction is contributed by both pH and movement or trafficking of the reporter. To investigate the impact of Spn-ATP reporter movement or trafficking on the determination of pH and ATP_spine,_ we performed the following calculations for the average fluorescence intensity of mOrg in Spn-ATP at each time point and treatment condition.

We determined the normalized change in intensity of EGFP from Homer2-EGFP-Luc, reflecting movement or trafficking, at each time point as:

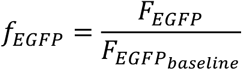

where *F*_EGFPbaseline_ is the average fluorescence intensity of EGFP at baseline.

We determined the normalized change in intensity of mOrg from Spn-ATP contributed by both pH changes and reporter movement or trafficking as:

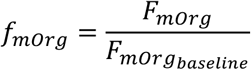

where *F_mOrg_*_baseline_ is the average fluorescence intensity of mOrg at baseline.

We determined the contribution of pH change alone to the mOrg fluorescence intensity at each time point as:

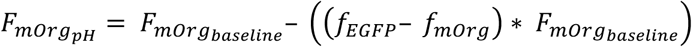

We used *F_mOrg_*_baseline_ to calculate the cytosolic pH at each time point as:

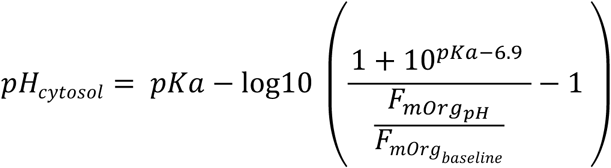

where 6.9 is the baseline cytosolic pH, pKa is 6.5 for mOrg.

We determined the contribution of movement or trafficking alone to the mOrg fluorescence intensity at each time point as:

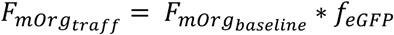

Since the *F_mOrg_*_traff_ was determined at cytosolic pH, 6.9, we calculated the corresponding fluorescence at pH 7 as the ATP calibration was done at pH 7.

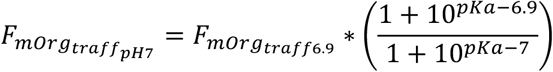

where pKa is 6.5 for mOrg.

*F_mOrgtraffpH7_* was used to normalize the movement or trafficking of Spn-ATP at all time points post-plasticity induction, and the L/F_traffcorr_ ratios were calculated as:

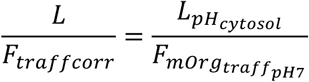

where *L*_pHcytosol_ is the average luminescence intensity at each time point at cytosolic pH.

The above L/F_traffcorr_ ratios were converted to [ATP], knowing the impact of pH on the K_m_ of ATP^6^ using a modified Michaelis-Menten equation as:

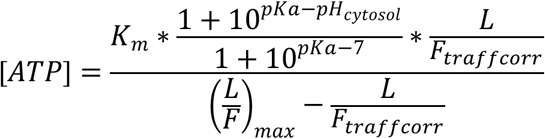

where the K_m_ and (L/F)_max_ or V_max_ are the Michaelis-Menten parameters obtained from the ATP titration curve measured at pH 7.0, and pK_a_ is pK_a_ of luciferase, 7.03.

### Spine structural plasticity measurements and analysis

Neurons were transfected with Spn-ATP and GCaMP6s or mEGFP and RCaMP plasmid constructs, along with MCU shRNA, when specified. Transfected neurons were identified by a change in GCaMP6s or RCaMP fluorescence corresponding to calcium transients in dendrites and spines. Spn-ATP or mEGFP fluorescence was used to identify spines for spine stimulation by two-photon glutamate uncaging. We did not differentiate between mushroom, stubby, or other spine shapes. Before glutamate uncaging, neurons were replaced with one μM TTX (Citrate salt, 2 mM stock made in water), 50 μM Forskolin (Tocris Bioscience, 25 mM stock made in DMSO), and 2 mM 4-Methoxy-7-nitroindolinyl-caged-L-glutamate (MNI caged glutamate) (Tocris Bioscience, 100 mM stock made in E4 buffer) in modified E4 buffer lacking Mg^2+^ (see above). Glutamate uncaging was performed using a multiphoton-laser 720 nm (Mai TAI HP) and a Pockel cell (ConOptics) to control the uncaging pulses. Spines at least 50 μm from the cell body were chosen for uncaging experiments (as we use dissociated cultures, we could not differentiate between apical and basal dendrites). We were blind to the mitochondrial compartments in the neurons, and the spines were chosen solely based on their response to an uncaging pulse. To test a spine’s response to an uncaging pulse, an uncaging spot (2 μm^2^) close to a spine head was selected, and two to three uncaging pulses at 10 ms pulse duration per pixel and 7.5-9.5 mW power were given and checked for spine-specific calcium transients. An uncaging protocol of 60 uncaging pulses at 0.5 Hz with 10 ms pulse duration per pixel at 7.5-9.5 mW power was used. For the CaMKII block, neurons were treated with 1 μM myrCN27 (1 μM stock made in DMSO) for 30 min. For Control vs. MCU KD experiments, images were acquired before plasticity induction at t = 0 min, at the end of the plasticity induction at t = 2 min, and then every 5 min for up to 30 min at t = 7, 12, 17, 22, 27 min. For Control vs. myrCN27 experiments, images were acquired 3 min before plasticity induction at t = −2, −1, 0 min; at the end of the plasticity induction at t = 2 min; and then every 1 min for up to 12 min.

For image analysis, GCaMP6s or mEGFP images were used. For spine-head width measurements using Spn-ATP and GCaMP6s, the GCaMP6s images from all time points (baseline: −2, −1 and 0 min, plasticity induction: 2 min, and post-plasticity induction: 2 to 12 mins) were concatenated and corrected for XY drifts using the StackReg plugin. For spine-head width measurements using mEGFP and RCaMP, ten images averaged from the pre-plasticity induction time point and one image from each post-plasticity induction time point were used. The background was subtracted using a rolling ball filter of a radius of 50 pixels using ImageJ. Next, a line crossing the center of the spine-head was drawn using a custom-written MATLAB script as before^2^. Then, the fluorescence intensity measured along the line was fit to a Gaussian to obtain the full width at half maxima (FWHM)–defined as the spine-head width^63^. As the FWHM is independent of fluorescence intensity, it was used to measure spine-head width even from fluctuating GCaMP6s fluorescence in spines.

For spine size and spine ATP correlation analysis, linear correlation fits were plotted for spine size vs. ATP_spine_, spine size vs. change in ATP_spine_ upon plasticity induction, ATP_spine_ vs. change in ATP_spine_ upon plasticity induction, and change in spine size vs. change in ATP_spine_ upon plasticity induction, resulting in Pearson’s r correlation coefficients.

### ATP_mito_ imaging and analysis

Neurons were transfected with Mito-ATP and GCaMP6s, along with the MCU shRNA, VAP sgRNA, or Cas9 control, when specified. Transfected neurons with mature spines were identified by a change in GCaMP6s fluorescence corresponding to calcium transients. In addition, the mCh signal from Mito-ATP was used to examine its expression and differentiate between dendritic (long ∼20-30 μm) and axonal (short 1-2 μm) mitochondria.

Before glutamate uncaging and imaging, the imaging buffer was replaced with 2 mM luciferin (Promega) for the bioluminescence reaction to detect ATP in the mitochondria, 1 μM TTX (Citrate salt, 2 mM stock made in water), 50 μM Forskolin (Tocris Bioscience, 25 mM stock made in DMSO), and 2 mM 4-Methoxy-7-nitroindolinyl-caged-L-glutamate (MNI caged glutamate) (Tocris Bioscience, 100 mM stock made in E4 buffer) in modified E4 buffer lacking Mg^2+^ (see above). Sample images of the selected neurons were acquired simultaneously in the 561 nm (mCh) and 488 nm (GCaMP6s) channels at exposure times of 50 ms each, and bioluminescence was acquired over 20 s. The neuron with the best mCh fluorescence was selected for further experiments. Dendritic branches at least 100 µm away from the soma, with mature mushroom-shaped spines near mitochondrial compartments, showing visible ATP signal within mitochondria, and are in the same focal plane as the spine, were chosen for two-photon glutamate uncaging. Glutamate uncaging was performed using a multiphoton-laser 720 nm (Chameleon, Coherent) and a Pockels cell (Conoptics) for controlling the uncaging pulses. To test a spine’s response to an uncaging pulse, an uncaging spot (2 μm^2^) close to a spine head was selected, and one to two uncaging pulses at 10 ms pulse duration per pixel and 7.5 mW back aperture power were given and checked for spine-specific calcium transients. For spine plasticity induction, an uncaging protocol of 60 uncaging pulses at 0.5 Hz with 10 ms pulse duration per pixel at 6 mW back aperture power of the 720 nm laser was used. An automated program was used for imaging the baseline (every 20 s for 3 min), two-photon glutamate uncaging to induce synaptic plasticity induction (2 min), and post-plasticity induction (every 20 s for 12 min) of the mitochondria of interest with simultaneous acquisitions in the 488 (GCaMP6s), 561 (mCh) and luminescence (luciferase bioluminescence) channels using the same exposure and acquisition times as mentioned above, using the 6D feature of the Slidebook 6 or 2024 software.

Images were analyzed using ImageJ to measure ATP at each time point. 2 µm length regions of interest were drawn on the mitochondrial compartments at the base of the plasticity-induced spine. Background subtracted luminescence intensity (L) from the mitochondrial region of interest was divided by the corresponding background subtracted mOrg fluorescence intensity (F) to determine the L/F ratio of the mitochondria of interest at each time point as:

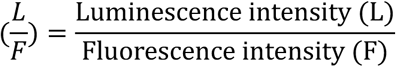

It was challenging to calibrate the Mito-ATP reporter, similar to the Spn-ATP reporter, as the permeabilization protocol required to obtain an ATP clamp was not consistently successful due to the mitochondrial double membrane beside the neuronal cell membrane. So, the calculated L/F values from Mito-ATP were interpreted as mitochondrial ATP measurements following pH correction (see below). For measuring the spatial profile of ATP, a custom-written script was used in ImageJ. The images from each time point of baseline and plasticity induction were stacked together and corrected for any drift in the XY plane using the StackReg plugin. 40 µm long regions of interest were drawn along the dendrites on either side of the plasticity-induced spine using the segmented line plugin, Ridge Detection. Negative distance values denote the direction of the cell body, whereas positive distance values denote the direction of the dendritic tip. The background-subtracted L and F intensities from these regions were calculated every 0.2 µm along the segmented line. The change in luminescence (ΔL) was calculated by subtracting the L at the 2 min post-plasticity induction time point from the corresponding average baseline L of the same dendritic region of interest. The ΔL along the dendrite was plotted at 5 µm intervals (with each data point obtained from a flanking average of ± 0.5 μm) to get the spatial profile of mitochondrial ATP increase.

### Mitochondrial pH imaging and pH correction for ATP_mito_

We initially attempted to perform pH corrections for Mito-ATP, similar to Spn-ATP, using the pH-sensitive mOrg fluorescent protein for pH-corrected, ratiometric luminescence to fluorescence (L/F) readouts (see above). However, mitochondrial pH determination using NH_4_Cl (pH 7.4) treatment was not reliable with mOrg, perhaps due to the alkaline baseline pH of mitochondria (pH 7.2, Fig. S3B)^17^, where the pKa of mOrg (6.5) was not sensitive enough. So, for ATP_mito_ we performed pH corrections by separate mitochondrial pH measurements using a pH-sensitive reporter targeted to the mitochondrial matrix, mito-pHluorin (pKa 7.1), as previously described^17,19^. Neurons were transfected with mitochondrial pHluorin (Mito-pHluorin) and RCaMP, along with MCU shRNA, VAP sgRNA, or Cas9 control when specified. The same imaging protocol as the ATP_mito_ measurements described above was used. Images were captured in the 488 channel for pHluorin, and 561 channel for RCaMP for baseline (3 min), 2-photon glutamate uncaging for plasticity induction (2 min), and post-plasticity induction (12 min). Following post-plasticity induction imaging, the E4 imaging buffer was aspirated and replaced with NH_4_Cl solution containing (in mM) 50 NH_4_Cl, 70 NaCl, 2.5 KCl, 2 CaCl_2_, 2 MgCl_2_, 30 HEPES (buffered to pH 7.4 at 37 °C) and 25 glucose, while timelapse images were continuously acquired every 15 s for 5 min and following NH_4_Cl addition.

Image analysis was performed using ImageJ as described above. The images from all time points (baseline, plasticity induction, post-plasticity induction, and NH_4_Cl treatment) in the pHluroin (488 nm) channel were stacked, and the StackReg plugin was used to correct for any drift in XY. 2 µm length regions of interest were drawn on the mitochondrial compartments at the base of the plasticity-induced spines as described for ATP_mito_ measurements. The background-subtracted fluorescence intensities of mito-pHluorin were calculated from these regions of interest at the respective time points. The mito-pHluorin fluorescence measured upon NH_4_Cl addition was called F_Max_ and the fluorescence at baseline was called F.

Mitochondrial pH of baseline and plasticity-induction were determined using the following modified Henderson-Hasselbalch equation^6^:

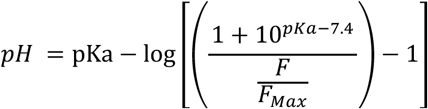

where pKa for MitopHluorin is 7.1^17^.

From the calculated pH values at various conditions, the L/F value corresponding to ATP_mito_ was pH corrected as follows:

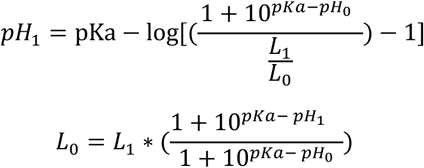

where pKa is 7.03 for ATP_mito_^6^, pH_0_ is the average pH at baseline, pH_1_ is the pH at the plasticity-induction and post-plasticity induction time points, L_0_ is the luminescence at pH_0,_ and L_1_ is the luminescence at pH_1_. The average pH at each condition was used for correction, and we did not propagate errors in pH measurements into the final error for L/F determination.

### Measurement of mitochondrial, ER, and spine calcium upon spine stimulation

To measure mitochondrial matrix and outer membrane (OMM) calcium, neurons were transfected with MatrixGCaMP or OMMGCaMP, respectively, together with RCaMP plasmid constructs. Transfected neurons and spines for stimulation were identified by changes in RCaMP fluorescence corresponding to calcium transients. To measure ER calcium, neurons were transfected with ERGCaMP210 and PSD95-mCherry. Transfected neurons were identified by ERGCaMP210 fluorescence, and the spines for stimulation were identified by PSD95-mCherry fluorescence. Before glutamate uncaging, the neuronal imaging buffer was replaced with 1 μM TTX (Citrate salt, 2 mM stock made in water), 2 mM 4-Methoxy-7-itroindolinyl-caged-L-glutamate (MNI caged glutamate) (Tocris Bioscience, 100 mM stock made in modified E4 buffer) in modified E4 buffer lacking Mg^2+^ (see above). Glutamate uncaging was performed using a multiphoton-laser 720 nm (Chameleon, Coherent) and a Pockels cell (Conoptics) to control the uncaging pulses. To test a spine’s response to an uncaging pulse, an uncaging spot (2 μm^2^) close to a spine-head was selected, and two to three uncaging pulses at 10 ms pulse duration per pixel and 7-8 mW back aperture power were given and checked for spine-specific calcium transients. For mitochondrial matrix or OMM calcium experiments, spines were stimulated with an uncaging protocol of 30 uncaging pulses at 0.3 Hz with 10 ms pulse duration per pixel at 7-8 mW power. However, for ER calcium experiments, the standard spine stimulation protocol (0.3 Hz 30 uncaging pulses with 10 ms pulse duration per pixel) did not induce observable ER calcium release^2^. So, we used a spine stimulation protocol of 6 uncaging pulses at 0.25 Hz with 100 ms pulse duration per pixel at 7-8 mW back aperture power. For the NMDAR block experiment, neurons were treated with 50 μM APV (Tocris, 100 mM stock made in modified E4 buffer lacking Mg^2+^) for at least 5 min. For the L-type calcium channel block experiment, neurons were treated with 10 μM Nimodipine (Tocris, 100 mM stock made in DMSO) for at least 10 min. For SERCA block experiments, neurons were treated with 1 μM Thapsigargin (Tocris, 1 mM stock made in DMSO) or 50 μM of CPA (Alomone, 50 mM stock made in DMSO) for at least 10 min. For RyR block experiments, neurons were treated with 100 μM Ryanodine (Tocris, 25 mM stock made in DMSO) for at least 5 min. For IP3R block experiments, neurons were treated with 3 μM XestC (Tocris, 100 μM stock made in DMSO) for at least 40 min.

Image analysis was done by ImageJ. ∼2 μm length regions of interest (ROI) were drawn at the base of the stimulated spine in the 488 channel to measure mitochondrial matrix, OMM, or ER calcium intensity (GCaMP), and in the 561 channel to measure dendritic calcium intensity (RCaMP). For spine calcium measurements, 1 μm diameter ROIs were drawn on the spine of interest and measured in the 561 channel (RCaMP). For mitochondrial matrix calcium experiments, only dendrites that showed mitochondrial matrix calcium response upon control spine stimulation were used for subsequent inhibitor treatments (using APV, Nimodipine, Thapsigargin, CPA, Ryanodine, and XestC) and further analyzed. For ER calcium experiments, only dendrites that showed ER calcium response upon control spine stimulation were used for subsequent inhibitor treatments (using Thapsigargin, CPA, Ryanodine, and XestC) and further analyzed. The average intensity from each ROI was measured for each channel and each time point. The average intensities were background subtracted using the intensity measured from an adjacent background area. For each successive time point during and after stimulation, the normalized intensity (ΔF/F) was calculated using the equation: ΔF/F = (F-F_0_)/F_0_, where F_0_ is defined as the average fluorescence intensity measured before spine stimulation, and F is defined as the fluorescence intensity measured at the time point of interest. Peak intensity values were identified using a Python script that detected local maxima. For the Control and MCU KD experiments, peak values from the ΔF/F time course were identified using the ‘find peaks’ function under the ‘peak analyzer’ tool in Origin with the ‘local maximum’ method. The baseline mode was set to ‘None (Y=0)’ and the direction of the peaks was specified as ‘positive’, and the number of local points was set to 1. The first ten peak intensity values of a time series were averaged to get the average ΔF/F corresponding to the spine of interest (ROI) following spine stimulation.

For mitochondrial matrix spatial profile measurements, images were averaged across the first 10 stimulation time points. A line of 1 μm thickness that is 20 μm equidistant on either side of the stimulated spine was drawn manually using the ‘segmented line’ tool along the dendrite. The stimulated spine position was marked as 0 μm. Negative distance values denote the direction of the cell body, whereas positive distance values denote the direction of the dendritic tip. The fluorescent profile of the line drawn along each dendrite was measured every 0.2 μm by the ‘plot profile’ function in ImageJ. The plot profile was background subtracted using the intensity measured from an adjacent background area. For each data point along the dendrite, the normalized intensity (ΔF/F) was calculated using the equation: ΔF/F = (F-F_0_)/F_0_, where F_0_ is defined as the average fluorescence intensity measured before spine stimulation, and F is defined as the fluorescence intensity measured poststimulation. The ΔF/F along the dendrite was plotted at 5 μm intervals (with each data point obtained from a flanking average of ± 2.4 μm) to get the spatial profile of mitochondrial matrix calcium increase.

### Correlation between dendritic and mitochondrial calcium

To determine the correlation between dendritic calcium and mitochondrial calcium uptake, neurons were transfected with MatrixGCaMP and RCaMP, respectively. A single uncaging pulse was given to a spine of interest, and the corresponding mitochondrial compartment at the base of the stimulated spine was imaged to monitor mitochondrial uptake. When the mitochondrial calcium uptake following spine stimulation cleared and decayed to baseline levels, the above stimulation protocol was repeated with a higher uncaging power, increasing by 0.6 mW 720 nm back aperture laser power for each condition, spanning 0 to 12 mW.

### Quantification and statistical analysis

Statistical details of experiments, including statistical tests and n and p values, are mentioned in the figure legends. Each animal corresponds to one weekly batch of neuronal culture preparation. In all the figures, the box whisker plots represent the median (line), mean (point), 25th-75th percentile (box), 10th-90th percentile (whisker), 1st–99th percentile (X), and min-max (_) ranges. Error bars in bar graphs and line traces are SEM. A p-value of less than or equal to 0.05 was considered significant for all statistical tests. Paired Sign Test and Mann-Whitney Test were used when datasets were rejected by the Shapiro-Wilk Normality Test. No statistical method was used to predetermine the sample size. Sample sizes were similar to or larger than those reported in the previous publications in the field and sufficient for our claims based on statistical significance.

