## Supplemental figures and legends, and video legends for "Synapses drive local mitochondrial ATP synthesis to fuel plasticity"

### Supplementary figures and legends

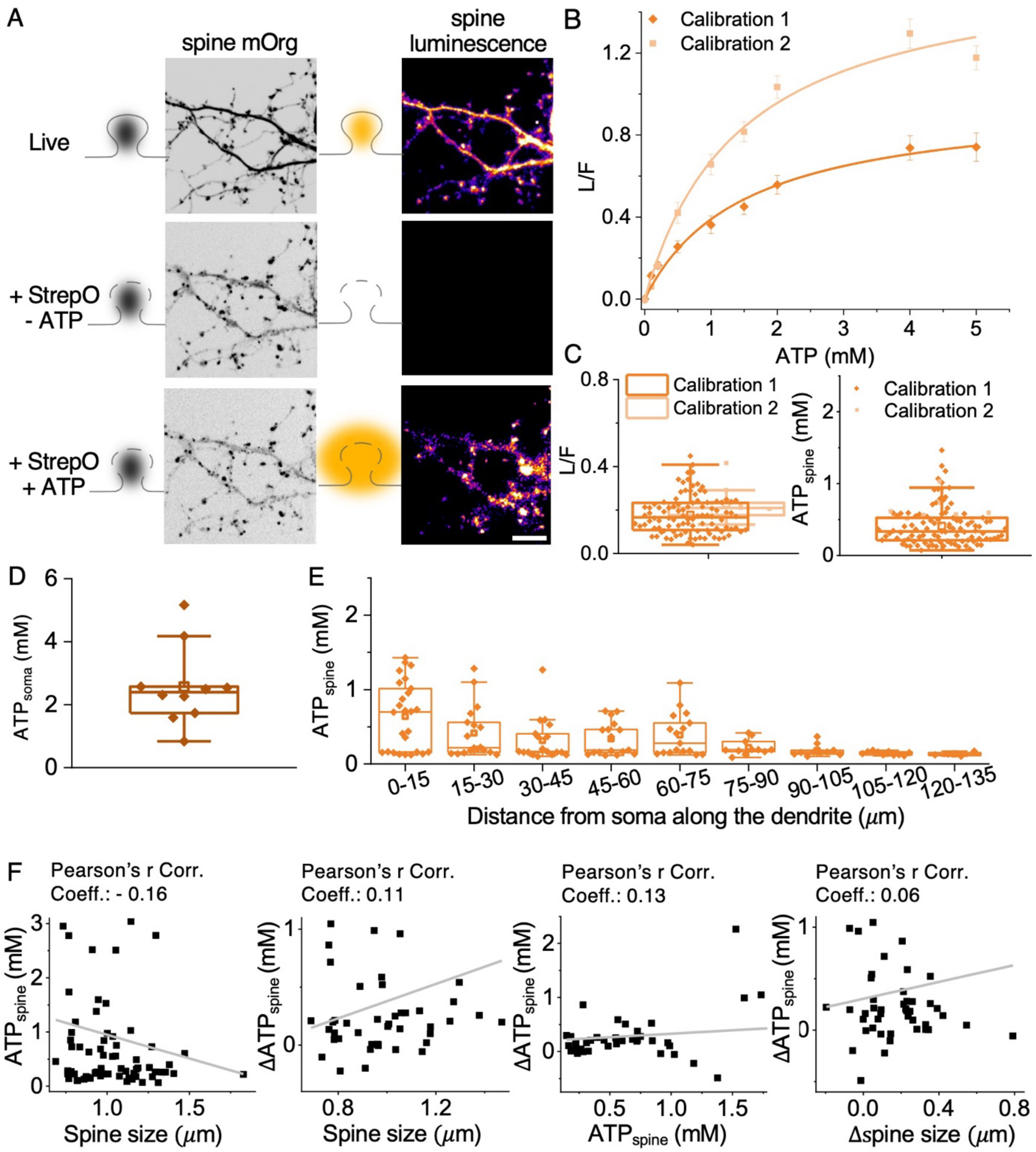


**Figure S1. Related to Figure 1. Control experiments for $ATP_{spine}$ measurements**

**A** Representative images (of **B**) of Spn-ATP expression (black, spine mOrg) and Spn-ATP luminescence (orange, spine luminescence) before (Live) and after permeabilization using Streptolysin-O (+StrepO, -ATP) show complete loss of spine luminescence upon permeabilization, while Spn-ATP expression is unaffected. The addition of a known concentration of ATP in the extracellular imaging buffer (+StrepO, +ATP) resulted in an ATP clamp across the permeabilized spine, and the corresponding luminescence signal for that ATP concentration was used for determining the ATP titration curve. **B** Michalis-Menten fit of the ATP titration curve obtained from two separate calibrations over the course of the project shows consistent kinetics, $K_m$ (Calibration 1)=1.5 $\pm$ 0.2 mM, $K_m$ (Calibration 2)=1.5 $\pm$ 0.3 mM, while the $V_{max}$ (i.e., $L/F_{max}$) changed, $V_{max}$ (Calibration 1)=0.97 $\pm$ 0.05, $V_{max}$ (Calibration 2)=1.65 $\pm$ 0.11, due to the change in laser power used for mOrg imaging over the course of the project (see Methods). n in spines, animals: 155, 8 (Calibration 1), 93, 3 (Calibration 2). **C** L/F and $ATP_{spine}$ measurements from individual spines in Fig 1B, C averaged across neurons show heterogeneity with a CV of 67%. n in neurons, animals: 109, 41. **D.** The average $ATP_{soma}$ measurements show a mean of 2.6 $\pm$ 0.4 mM. n in soma, animals: 10, 5. **E** The average $ATP_{spine}$ measured in spines at increasing distances from the soma decreases along the dendrite. n in spines, animals: 26, 5 (0-15 μm), 18, 3 (15-30 μm), 19, 4 (30-45 μm), 17, 5 (45-60 μm), 17, 4 (60-75 μm), 11, 5 (75-90 μm), 10, 3 (90-105 μm), 10, 2 (105-120 μm), 4, 2 (120-135 μm). **F.** Linear fit between spine size and $ATP_{spine}$; spine size and $ATP_{spine}$ change ($\Delta ATP_{spine}$); $ATP_{spine}$ and $ATP_{spine}$ change ($\Delta ATP_{spine}$); and spine size change ($\Delta$spine size) and $ATP_{spine}$ change ($\Delta ATP_{spine}$) do not show strong correlations. $ATP_{spine}$ vs. spine size, Linear fit, Intercept: 1.83 $\pm$ 0.71, Slope: -0.88 $\pm$ 0.67, p-value (slope): 0.19602. $\Delta ATP_{spine}$ vs. Spine size, Linear fit, Intercept: -0.36 $\pm$ 1.07, Slope: 0.74 $\pm$ 1.07, p-value (slope): 0.49; $\Delta ATP_{spine}$ vs. $ATP_{spine}$, Linear fit, Intercept: 0.2 $\pm$ 0.27, Slope = 0.13 $\pm$ 0.15, p-value (slope): 0.39; $\Delta$ATP vs. $\Delta$spine size, Linear fit, Intercept: 0.30 $\pm$ 0.25, Slope = 0.41 $\pm$ 1.06, p-value (slope): 0.70.

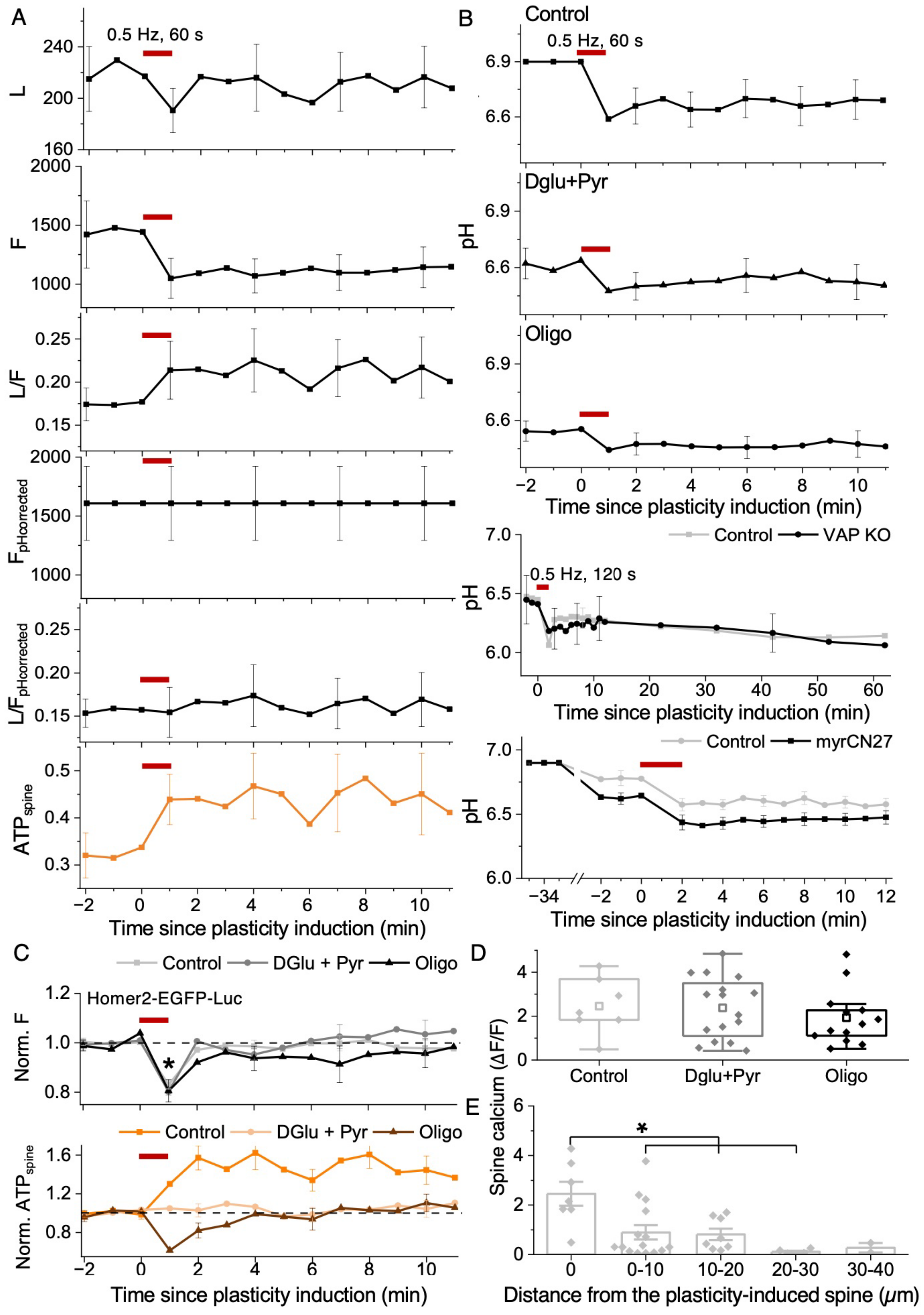

A
0.5 Hz, 60 s
L
F
L/F
F pHcorrected
L/F pHcorrected
ATP spine
Time since plasticity induction (min)
B
Control
Dglu+Pyr
Oligo
pH
Time since plasticity induction (min)
Control
VAP KO
0.5 Hz, 120 s
Control
myrCN27
C
Control
DGlu + Pyr
Oligo
Homer2-EGFP-Luc
Norm. F
Norm. ATP spine
D
Spine calcium (ΔF/F)
Control
Dglu+Pyr
Oligo
E
Distance from the plasticity-induced spine (μm)

**Figure S2. Related to Figure 2. pH and movement corrections for $ATP_{spine}$ and spine calcium measurements**

**A** The average time course of background subtracted L, F, L/F, $F_{pHcorrected}$, and $L/F_{pHcorrected}$ used for $ATP_{spine}$ calculations upon synaptic plasticity induction (red bar, 0.5 Hz, 60 s) (see Methods). n in spines, animals: 10, 3. **B** The average time course of pH changes upon synaptic plasticity induction (red bar, 0.5 Hz, 60 or 120 s) in Control, DGlu+Pyr, Oligo, Cas9 Control, VAP KO, and myrCN27-treated neurons and its Control used for $ATP_{spine}$ pH corrections (see Methods). n in spines, animals: 10, 3 (Control), 13, 3 (Oligo), 18, 3 (Dglu+Pyr), 10, 7 (Cas9 Control), 8, 7 (VAP KO), 16, 3 (Control for myrCN27), 23, 7 (myrCN27). **C** The average time course of normalized background subtracted EGFP fluorescence (F) from a modified Spn-ATP reporter, Homer2-EGFP-Luc, where mOrg was replaced with EGFP to eliminate its pH sensitivity, showed a small but significant decrease upon spine plasticity induction (red bar, 0.5 Hz, 60 s) and recovery to baseline, indicating the reporter's trafficking or movement. $ATP_{spine}$ measurements corrected for the trafficking or movement of the Spn-ATP reporter gave similar results as in Fig. 2C (see Methods). n in spines, animals: 16, 3 (Control), 8, 3 (DGlu+Pyr), 6, 3 (Oligo)**.** Paired Sample t test, p-values: <0.0001 (Control: Baseline vs. 1 min post-plasticity induction), 0.27677 (Control: Baseline vs. 11 min post-plasticity induction), 0.00015 (DGlu+Pyr: Baseline vs. 1 min post-plasticity induction), 0.35196 (DGlu+Pyr: Baseline vs. 11 min post-plasticity induction), 0.010 (Oligo: Baseline vs. 1 min post-plasticity induction), 0.70283 (Oligo: Baseline vs. 11 min post-plasticity induction). **D** Spine calcium response ($\Delta F/F$) upon spine stimulation was unaffected in DGlu+Pyr (dark gray) and Oligo (black) treated spines compared to the Control (light gray). n in spines, animals: 7, 3 (Control), 16, 3 (Dglu+Pyr), 13, 3 (Oligo). One-way ANOVA, Tukey test, p-values: 0.99056 (Control vs. DGlu+Pyr), 0.67582 (Control vs. Oligo). **E** Spine calcium response ($\Delta F/F$) upon synaptic plasticity induction is significantly reduced in spines 0-30 $\mu$m from the plasticity-induced spine compared to the plasticity-induced spine at 0 $\mu$m. n in spines, animals: 7, 3 (0 $\mu$m), 15, 4 (0-10 $\mu$m), 8, 3 (10-20 $\mu$m), 5, 3 (20-30 $\mu$m), 2, 1 (30-40 $\mu$m). One-way ANOVA, Tukey test, p-values: 0.01147 (0 vs 0-10 $\mu$m), 0.02167 (0 vs. 10-20 $\mu$m), 0.00227 (0 vs. 20-30 $\mu$m), 0.06292 (0 vs. 30-40 $\mu$m).

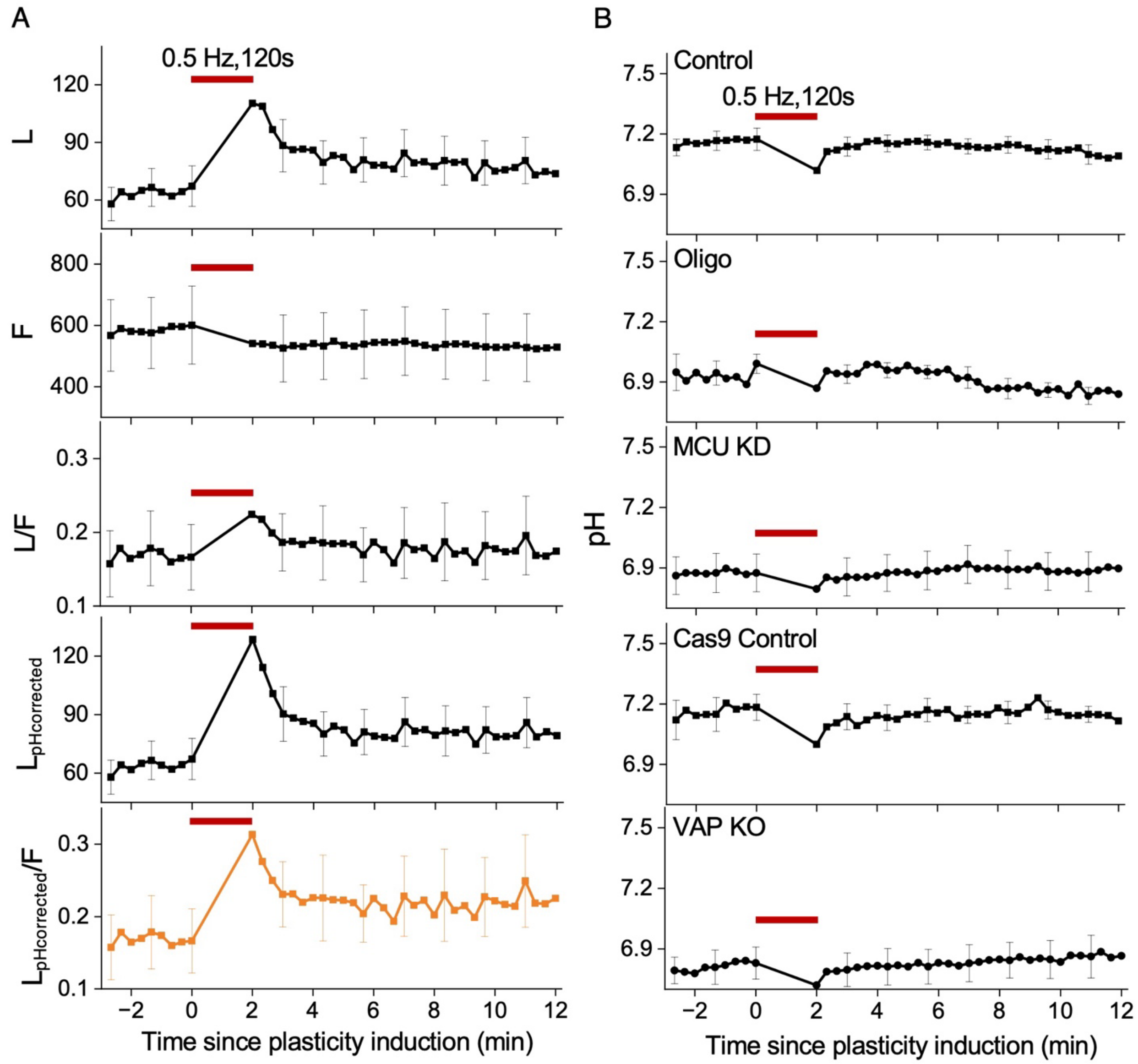


**Figure S3. Related to Figure 3. pH corrections for $ATP_{mito}$ upon synaptic plasticity induction**

**A** The average time course of background subtracted L, F, L/F, $L_{pHcorrected}$, and $L_{pHcorrected}$/F used for $ATP_{mito}$ calculations (see Methods) upon synaptic plasticity induction (red bar, 0.5 Hz, 120 s) shows an instant and sustained increase in the luminescence channel (black, L, L/F, orange, $L_{pHcorrected}$, $L_{pHcorrected}$/F) but not in the fluorescence channel (black, F). n in dendrites, animals: 14, 5. **B** The average time course of pH changes upon synaptic plasticity induction (red bar, 0.5 Hz, 120 s) in Control, Oligo, MCU KD, Cas9 Control, and VAP KO neurons used for mitochondrial ATP pH corrections (see Methods). n in dendrites, animals: 12, 4 (Control), 4, 2 (Oligo), 10, 4 (MCU KD), 9, 2 (Cas9 Control), 10, 2 (VAP KO).

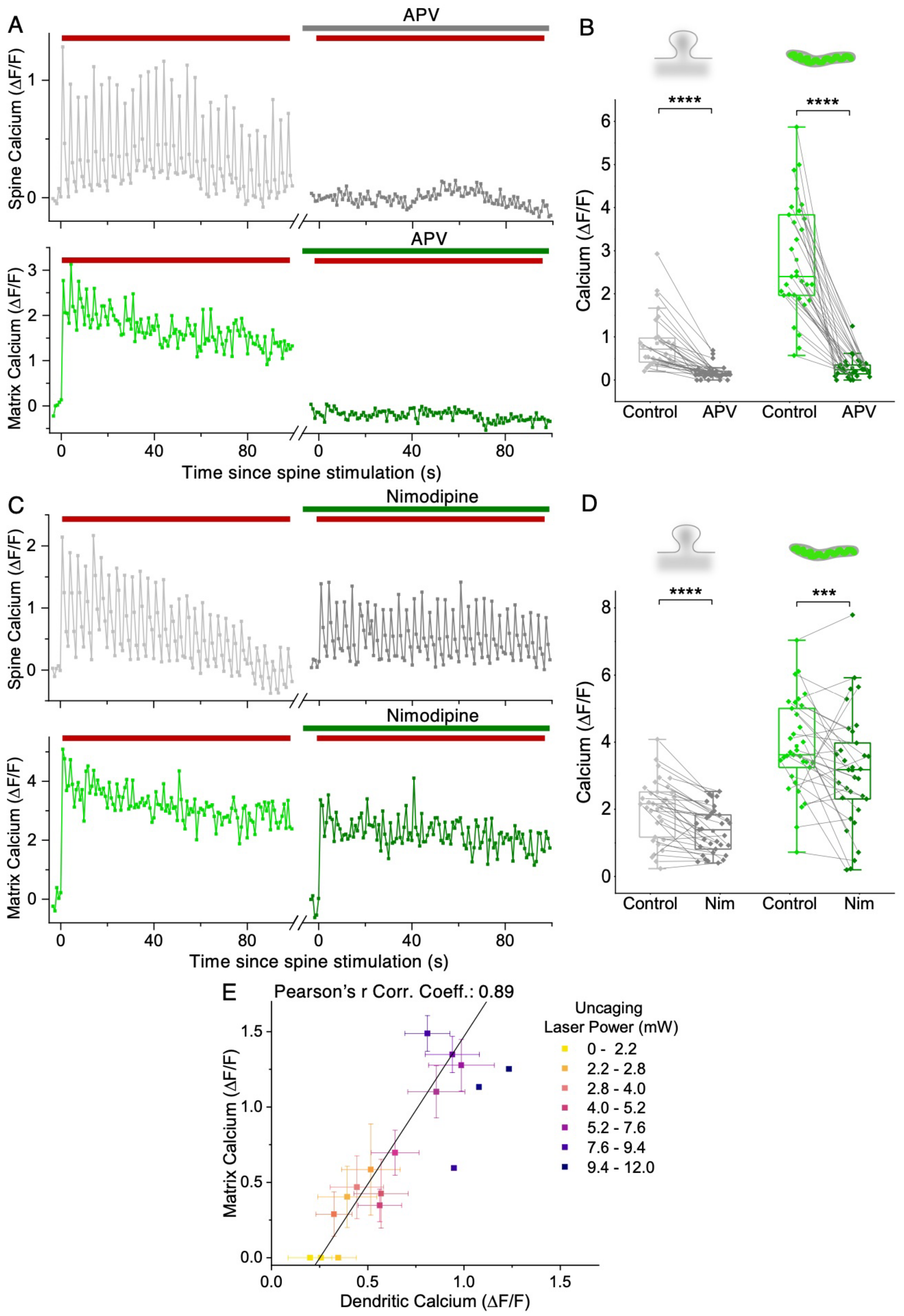

A
APV
Spine Calcium (ΔF/F)
Matrix Calcium (ΔF/F)
Time since spine stimulation (s)
B
Calcium (ΔF/F)
****
Control
APV
C
Nimodipine
D
***
Nim
E
Pearson's r Corr. Coeff.: 0.89
Matrix Calcium (ΔF/F)
Dendritic Calcium (ΔF/F)
Uncaging
Laser Power (mW)
0 - 2.2
2.2 - 2.8
2.8 - 4.0
4.0 - 5.2
5.2 - 7.6
7.6 - 9.4
9.4 - 12.0

**Figure S4. Related to Figure 4. Mitochondrial calcium entry is NMDAR-dependent, L-type calcium channel independent, and is linearly proportional to dendritic calcium**

**A** Representative time course (of **B**) of the spine (gray) and mitochondrial matrix (green) calcium responses ($\Delta F/F$) upon spine stimulation (red bar, 0.3 Hz 30 pulses) shows abolished spine calcium (dark gray) and mitochondrial calcium (dark green) upon NMDAR inhibition (using APV). **B** Spine and mitochondrial calcium uptake ($\Delta F/F$) upon spine stimulation is abolished upon NMDAR inhibition (using APV). n in dendrites, animals: 31, 3. Paired Sample Sign Test, p-value: <0.0001 (Spine calcium: Control vs. APV). Paired Sample t Test, p-value: <0.0001 (Matrix calcium: Control vs. APV). **C** Representative time course (of **D**) of the spine (gray) and mitochondrial matrix (green) calcium responses ($\Delta F/F$) during spine stimulation (red bar, 0.3 Hz 30 pulses) shows a significant but slight decrease in spine calcium (dark gray) and mitochondrial calcium (dark green) upon L-type calcium channel inhibition (using Nimodipine). **D** Spine and mitochondrial calcium uptake ($\Delta F/F$) upon spine stimulation shows a small but significant decrease upon L-type calcium channel inhibition (using Nimodipine). n in dendrites, animals: 33, 3. Paired Sample t Test, p-value: <0.0001 (Spine calcium: Control vs. Nimodipine). Paired Sample t Test, p-value: <0.00732 (Matrix calcium: Control vs. Nimodipine). **E** Mitochondrial Matrix and dendritic calcium response ($\Delta F/F$) upon spine stimulation are linearly correlated. Linear fit, Intercept: $-0.5 \pm 0.2$, Slope = $1.96 \pm 0.26$, p-value (slope): <0.0001. n in dendrites, animals: 31, 3.

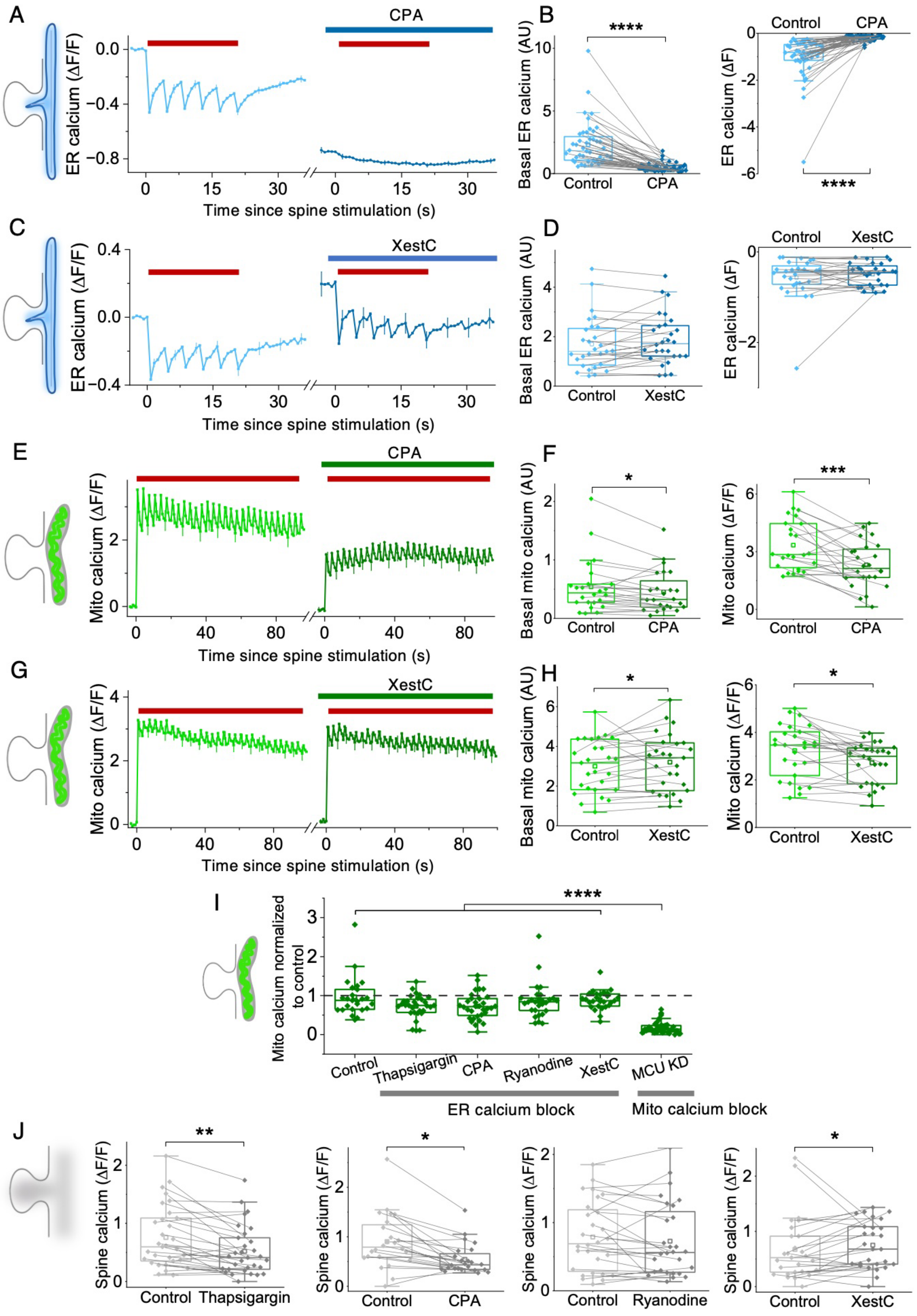

A
CPA
ER calcium (ΔF/F)
Time since spine stimulation (s)
B
Basal ER calcium (AU)
Control
CPA
****
ER calcium (ΔF)
C
XestC
D
E
Mito calcium (ΔF/F)
F
Basal mito calcium (AU)
*
***
G
H
I
Mito calcium normalized to control
Thapsigargin
Ryanodine
MCU KD
ER calcium block
Mito calcium block
J
Spine calcium (ΔF/F)
**

**Figure S5. Related to Figure 5. Control experiments for ER and mitochondrial calcium measurements**

**A, C** The average time course of ER calcium response (ΔF/F) during spine stimulation (red bar, 0.2 Hz, 6 pulses) shows depletion of ER calcium stores upon CPA but not upon XestC treatment. n in dendrites, animals: 44, 3 (CPA), 28, 3 (XestC). **B, D** Basal ER calcium and ER calcium release (ΔF) upon spine stimulation are significantly reduced upon CPA but not XestC treatment. n in dendrites, animals: 44, 3 (CPA), 28, 3 (XestC). Paired Sample Sign Test, p-values: <0.0001 (Basal ER calcium and ER calcium response: Control vs. CPA), 0.85011 (Basal ER calcium: Control vs. XestC), 0.85011 (ER calcium response: Control vs. XestC). **E, G** The average time course of mitochondrial calcium response (ΔF/F) during spine stimulation (red bar, 0.3 Hz, 30 pulses) shows a small decrease upon CPA and XestC treatments. n in dendrites, animals: 24, 3 (CPA), 27, 3 (XestC). **F, H** Basal mitochondrial calcium and mitochondrial calcium response (ΔF/F) upon spine stimulation show a small but significant decrease upon CPA and XestC treatments. n in dendrites, animals: 24, 3 (CPA), 27, 3 (XestC). Paired Sample t Test, p-values: 0.04934 (Basal mito calcium: Control vs. CPA), 0.00036 (Mito calcium response: Control vs. CPA). Paired Sample Sign Test, p-values: 0.02092 (Basal mito calcium: Control vs. XestC, 0.05429 (Mito calcium response: Control vs. XestC). **I** Mitochondrial calcium response (ΔF/F) during spine stimulation upon various treatments normalized to their respective untreated controls (DMSO) was significantly reduced when mitochondria calcium entry was blocked (using MCU KD) but not when ER calcium entry or release was blocked (using Thapsigargin CPA, Ryanodine, and XestC). n in dendrites, animals: 25, 3 (Control), 28, 3 (Thapsigargin), 24, 3 (CPA), 31, 3 (Ryanodine), 27, 3 (XestC), 44, 4 (MCU KD). One-way ANOVA, Tukey Test, p-values: 0.06844 (Control vs. Thapsigargin), 0.07631 (Control vs. CPA), 0.88466 (Control vs. Ryanodine), 0.96982 (Control vs. XestC), <0.0001 (Control vs. MCU KD). **J** Spine calcium response (ΔF/F) upon spine stimulation shows a small but significant reduction in Thapsigargin, CPA, and XestC, but not upon Ryanodine treatment. n in dendrites, animals: 31, 3 (Thapsigargin), 24, 3 (CPA), 28, 3 (Ryanodine), 27, 3 (XestC). Paired Sample Sign Test, p-values: 0.00406 (Control vs. Thapsigargin), 0.05429 (Control vs. XestC). Paired Sample t Test, p-values: <0.0001 (Control vs. CPA), 0.28639 (Control vs. Ryanodine).

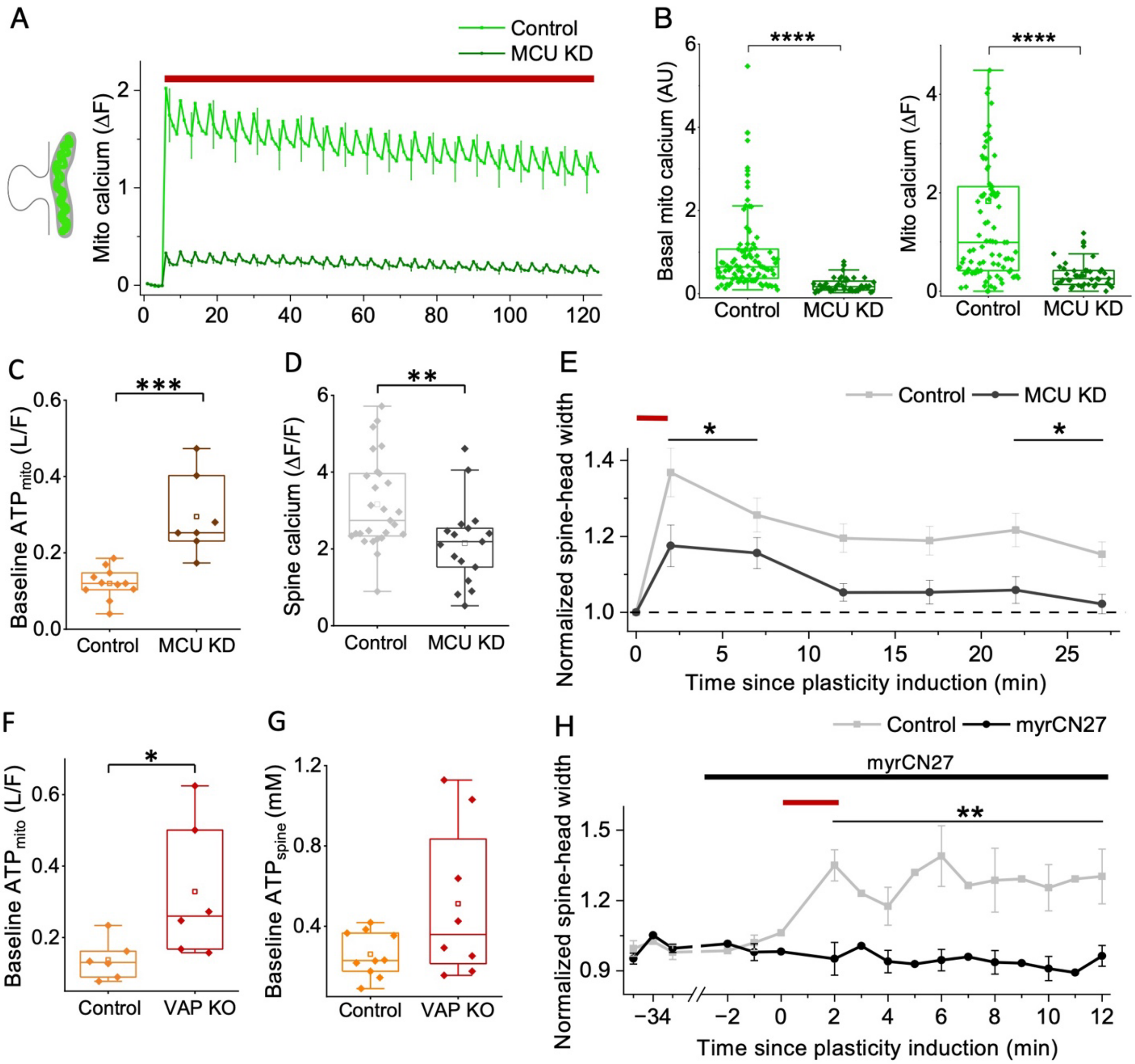


**Figure S6. Related to Figure 6. Control experiments for MCU, VAP, and CaMKII inhibition**

**A** The average time course of mitochondrial calcium response (green, ΔF/F) during spine stimulation shows a significant decrease in MCU KD (dark green) compared to the Control dendrites. n in dendrites, animals: 93, 4 (Control), 44, 4 (MCU KD). **B** Basal mitochondrial calcium and mitochondrial calcium response (ΔF/F) (green) upon spine stimulation were significantly reduced in MCU KD (dark green) compared to the Control dendrites. n in dendrites, animals: 93, 4 (Control), 44, 4 (MCU KD). Mann-Whitney Test, p-value: <0.0001 (Basal mito calcium and Mito calcium response: Control vs. MCU KD). **C** Basal ATP$_{mito}$ (L/F) shows a significant increase in MCU KD (brown) neurons compared to the Control (orange). n in dendrites, animals: 11, 6 (Control), 7, 3 (MCU KD). Two Sample t Test, p-value: 0.0001 (Control vs. MCU KD). **D** Spine calcium response ($\Delta F/F$) upon spine stimulation is significantly reduced in MCU KD (dark gray) compared to the Control (light gray). n in spines, animals: 27, 10 (Control), 17, 5 (MCU KD). Two-sample t-test, p-value: 0.0058 (Control vs. MCU KD). **E** The average time course shows a sustained rise in spine-head width in the Control (light gray) but not in MCU KD (dark gray) at 27 min post-plasticity induction (red bar, 0.5 Hz, 120 s). n in spines, animals: 27, 10 (Control), 17, 5 (MCU KD). One-way ANOVA, Tukey test, p-values: 0.01494 (2-7 min post-plasticity induction: Control vs. MCU KD), 0.01753 (22–27 min post-plasticity induction: Control vs. MCU KD). **F** Basal $ATP_{mito}$ (L/F) shows a significant increase in VAP KO (red) compared to the Control (orange). n in dendrites, animals: 6, 3 (Control), 6, 5 (VAP KO). Two Sample t Test, p-value: 0.04004 (Control vs. VAP KO). **G** Basal $ATP_{spine}$ was not statistically significant between VAP KO (red) and the Control (orange). n in spines, animals: 10, 7 (Control), 8, 7 (VAP KO). Two Sample t Test, p-value: 0.06456 (Control vs. VAP KO). **H** The average time course shows a sustained rise in spine-head width upon synaptic plasticity induction (red bar, 0.5 Hz, 120 s) in the Control (light gray) but not in myrCN27-treated neurons (black) post-plasticity induction 2-12 min. n in spines, animals: 12, 3 (Control), 11, 5 (myrCN27). One-way ANOVA, Tukey test, p-value: 0.010190 (2–12 min post-plasticity induction: Control vs. myrCN27).

**Supporting video legends**

**Supp. video 1. Instant and sustained increase in $ATP_{spine}$ upon synaptic plasticity induction**

Time-lapse videos of the representative images in Fig. 2B show an increase in spine calcium (GCaMP6s, white) and Spn-ATP luminescence (orange, luc) upon spine plasticity induction (white and black asterisk) without a change in Spn-ATP expression (black, mOrg). Each frame of the time-lapse represents 1 min. Frames 1 and 2: baseline -2 to 0 min, Frame 3: post-plasticity induction 1 min, Frames 4 - 6: post-plasticity induction 2 to 4 min. Scale bar 5 μm.

**Supp. video 2. Instant, local, and sustained increase in $ATP_{mito}$ upon synaptic plasticity induction**

Time-lapse videos of the representative images in Fig. 3B show an instant increase in spine calcium (GCaMP6s, white) and Mito-ATP luminescence (orange, Luc) upon spine plasticity induction (white and black asterisk) without a change in Mito-ATP expression (black, mCh). Each frame of the time-lapse represents 20 s. Frames 1 - 3: baseline -60 to 0 s, Frame 4: post-plasticity induction 20 s, Frames 5 - 7: post-plasticity induction 40 – 80 s. Scale bar 10 μm.

**Supp. video 3. Instant, local, and sustained increase in mitochondrial calcium upon spine stimulation**

**A** Time-lapse videos of the representative images in Fig. 3A show an increase in spine calcium (white, RCaMP) upon single spine stimulation (white asterisk), followed by an instant and spatially restricted increase in mitochondrial matrix calcium (green, MatrixGCaMP) at the base of the stimulated spine. Each frame represents 0.83 s. Stimulation happens every 4 frames, starting from frame 5. Scale bar 5 μm. **B** Time-lapse videos of the representative images in Fig. 3B show an increase in spine calcium (white, RCaMP) upon single spine stimulation (white asterisk), followed by a diffused increase in mitochondrial outer membrane calcium (green, OMMGCaMP) at the base of the stimulated spine. Each frame represents 0.83 s. Stimulation happens every 4 frames, starting from frame 5. Scale bar 5 μm.
